# Low Abundance of Circulating Tumor DNA in Localized Prostate Cancer

**DOI:** 10.1101/655506

**Authors:** S. Thomas Hennigan, Shana Y. Trostel, Nicholas T. Terrigino, Olga S. Voznesensky, Rachel J. Schaefer, Nichelle C. Whitlock, Scott Wilkinson, Nicole V. Carrabba, Rayann Atway, Steven Shema, Ross Lake, Amalia R. Sweet, David J. Einstein, Fatima Karzai, James L. Gulley, Peter Chang, Glenn J. Bubley, Steven P. Balk, Huihui Ye, Adam G. Sowalsky

**Affiliations:** Laboratory of Genitourinary Cancer Pathogenesis, National Cancer Institute, National Institutes of Health (NIH), Bethesda, MD, 20892; Department of Medicine, Beth Israel Deaconess Medical Center, Boston, MA, 02215; CCR Genomics Core, National Cancer Institute, NIH, Bethesda, MD, 20892; Genitourinary Malignancies Branch, National Cancer Institute, NIH, Bethesda, MD, 20892; Department of Surgery, Beth Israel Deaconess Medical Center, Boston, MA, 02215; Department of Pathology, Beth Israel Deaconess Medical Center, Boston, MA, 02215

**Keywords:** prostate cancer, circulating tumor DNA, biomarker, cell-free DNA

## Abstract

Despite decreased screening-based detection of clinically insignificant tumors, most diagnosed prostate cancers are still indolent, indicating a need for better strategies for detection of clinically significant disease prior to treatment. We hypothesized that patients with detectable circulating tumor DNA (ctDNA) were more likely to harbor aggressive disease. We applied ultra-low pass whole genome sequencing to profile cell-free DNA from 112 patients diagnosed with localized prostate cancer and performed targeted resequencing of plasma DNA for somatic mutations previously identified in matched solid tumor in nine cases. We also performed similar analyses on patients with metastatic prostate cancer. In all cases of localized disease, even in clinically high-risk patients who subsequently recurred, we did not detect ctDNA by either method in plasma acquired before surgery or before recurrence. In contrast, ctDNA was detected from patients with metastatic disease. Our findings demonstrate clear differences between localized and advanced prostate cancer with respect to the dissemination and detectability of ctDNA. Because allele-specific alterations in ctDNA are below the threshold for detection in localized prostate cancer, other approaches to identify cell-free nucleic acids of tumor origin may demonstrate better specificity for aggressive disease.

## INTRODUCTION

Over the last two decades, prostate cancer remains the most diagnosed neoplasm in American men, representing approximately 20% of all new diagnoses in 2019 [1]. Overtreatment of newly-diagnosed, indolent prostate cancers detected by rising levels of prostate-specific antigen (PSA) has been mitigated by increasingly widespread adoption of active surveillance, MRI-targeted biopsies, nomograms, and molecular tests for assessing the risk posed by unsampled higher grade disease [2–5]. While the absence of adverse pathological features, such as high Gleason score or seminal vesicle invasion, from biopsy is associated with improved outcomes following definitive therapy (surgery or radiation), sampling errors may lead to underestimation of the risk of biochemical recurrence. The potential for failure to detect pathologic features motivates increased biopsy frequency and premature withdrawal from active surveillance [6–8].

Numerous recent studies have explored the genomic basis for development of localized prostate cancer, showing distinct evolutionary paths in nonindolent versus indolent disease. The fate of tumors to progress from their somatic progenitors is set early, with alterations in *ATM*, *PTEN*, and *MYC* subclones having predictive power for the existence of higher grade disease including occult oligometastases at the time of radical prostatectomy [9–13]. As the vast majority of these alterations occur as copy number gains or deletions, the percentage of the genome affected by large chromosomal rearrangements is similarly predictive of biochemical recurrence and poor outcome [10, 14, 15].

Analysis of plasma cell-free DNA (cfDNA) has rapidly gained traction for profiling tumor genomics in patients with metastatic disease, especially in prostate cancer in which dissemination to the bone occurs frequently [16]. Allele-specific assays that detect major driver events, such as mutations to *AR*, *APC*, *EGFR*, and *ERBB2* are commercially-available for identification of recurrent, targetable clonal alterations in advanced stages of several cancers, including prostate, colorectal, lung, and breast cancer [17]. Comprehensive cancer panels, as well as whole genome and exome sequencing, can also be used to interrogate somatic copy number alterations (SCNAs) from plasma DNA, with varying resolution depending on the sequence modality and depth [18, 19]. Personalized, bespoke sequencing assays have shown sensitivity for the detection of urothelial and colorectal cancers [20, 21]. The success of these approaches has been thought to depend upon high tumor burden and the propensity of the tumor to shed circulating tumor DNA (ctDNA) into the bloodstream with proportional contribution of subclones to the ctDNA pool [22, 23]. However, the feasibility of applying these approaches to assess the clinical trajectory of newly diagnosed prostate cancer patients has not been established.

In this study, we performed ultra-low pass (ULP) whole genome sequencing (WGS) of cfDNA from 112 patients with localized prostate cancer to assess genome-wide SCNAs and their association with biochemical recurrence-free survival (median follow-up of 50 months). We also performed deeper, targeted sequencing of cfDNA in nine cases with matched multi-region sequencing of prostate tumor tissue to identify subclones in ctDNA that may associate with adverse pathologic features or mediate relapse. The absence of signal from ctDNA in plasma from localized, but not metastatic, prostate cancer patients, demonstrates that the strategy of using tumor-specific somatic alterations for assessing disease burden is of minimal clinical utility.

## RESULTS

### Large SCNA events are not detectable in the plasma of patients with localized prostate cancer

ULP-WGS has been proposed as a screening technique to detect large SCNAs in cfDNA for the rapid and inexpensive determination of ctDNA content [19]. To assess the feasibility of this analysis in patients with localized disease, blood was obtained from 112 consecutive patients (L001-L112) between April 2014 and January 2016. Patients consented to participate in tissue and blood procurement protocols while undergoing radical prostatectomy as definitive therapy for newly diagnosed prostate cancer or previously diagnosed prostate cancer that had progressed on active surveillance. Clinical demographics for this cohort are given in Table 1. Blood was collected from an additional seven consecutive patients (M01-M07) with radiographically-confirmed metastatic prostate cancer who would be expected to harbor ctDNA based on high tumor volumes (Table 1).

**Table 1.**
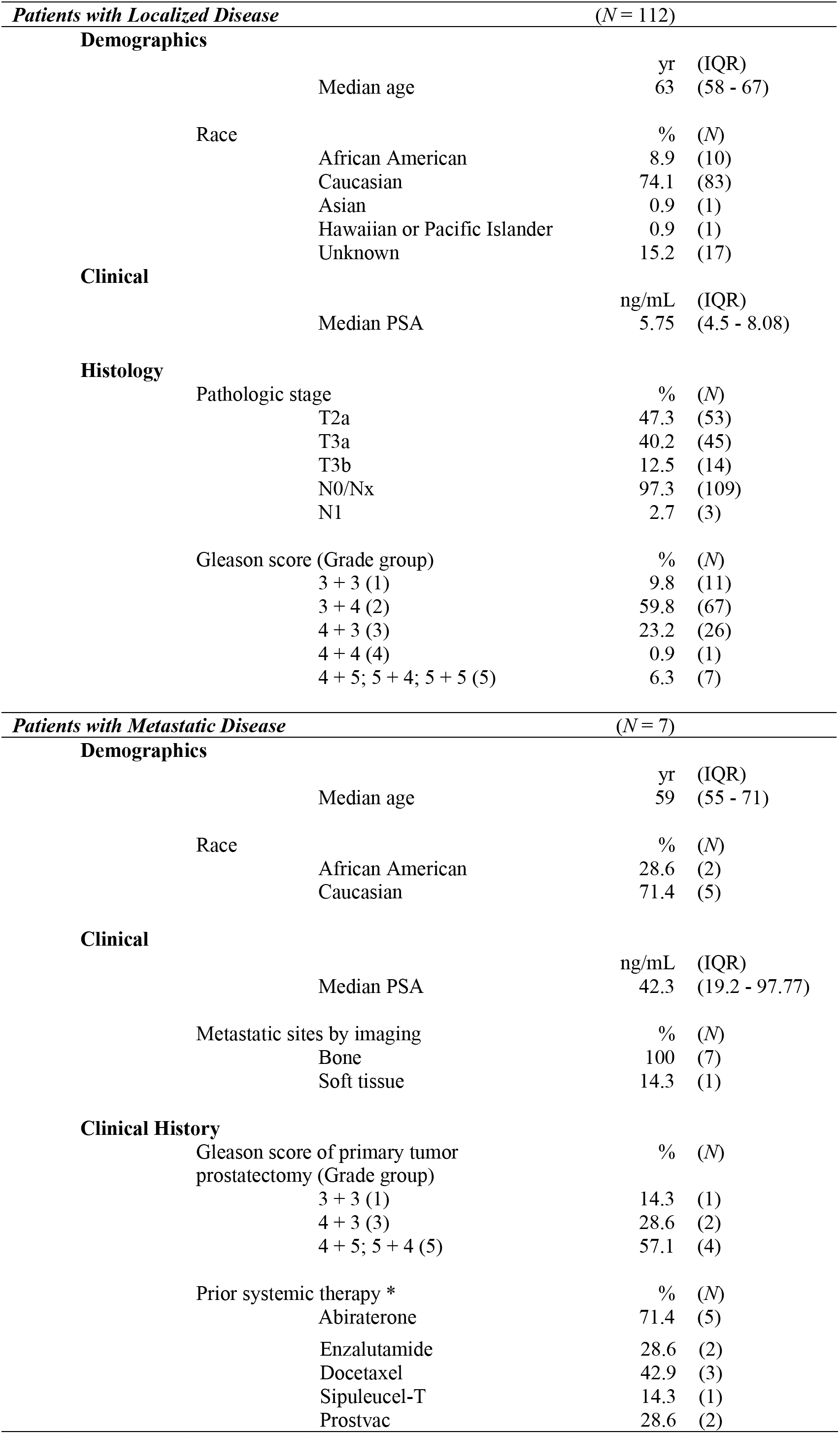
Patient characteristics for 112 men with localized prostate cancer and seven men with metastatic prostate cancer. IQR = interquartile range. Data are presented as *N* (IQR) or % (*N*). *Sum of percentages exceeds 100% due to patients receiving more than one prior therapy.

| *Patients with Localized Disease* |  | *(N = 112)* |  |
| --- | --- | --- | --- |
| **Demographics** |  |  |  |
|  | Median age | yr | (IQR) |
|  |  | 63 | (58 - 67) |
|  | Race | % | (N) |
|  | African American | 8.9 | (10) |
|  | Caucasian | 74.1 | (83) |
|  | Asian | 0.9 | (1) |
|  | Hawaiian or Pacific Islander | 0.9 | (1) |
|  | Unknown | 15.2 | (17) |
| **Clinical** |  |  |  |
|  | Median PSA | ng/mL | (IQR) |
|  |  | 5.75 | (4.5 - 8.08) |
| **Histology** |  |  |  |
|  | Pathologic stage | % | (N) |
|  | T2a | 47.3 | (53) |
|  | T3a | 40.2 | (45) |
|  | T3b | 12.5 | (14) |
|  | N0/Nx | 97.3 | (109) |
|  | N1 | 2.7 | (3) |
|  | Gleason score (Grade group) | % | (N) |
|  | 3 + 3 (1) | 9.8 | (11) |
|  | 3 + 4 (2) | 59.8 | (67) |
|  | 4 + 3 (3) | 23.2 | (26) |
|  | 4 + 4 (4) | 0.9 | (1) |
|  | 4 + 5; 5 + 4; 5 + 5 (5) | 6.3 | (7) |

| *Patients with Metastatic Disease* |  | *(N = 7)* |  |
| --- | --- | --- | --- |
| **Demographics** |  |  |  |
|  | Median age | yr | (IQR) |
|  |  | 59 | (55 - 71) |
|  | Race | % | (N) |
|  | African American | 28.6 | (2) |
|  | Caucasian | 71.4 | (5) |
| **Clinical** |  |  |  |
|  | Median PSA | ng/mL | (IQR) |
|  |  | 42.3 | (19.2 - 97.77) |
|  | Metastatic sites by imaging | % | (N) |
|  | Bone | 100 | (7) |
|  | Soft tissue | 14.3 | (1) |
| **Clinical History** |  |  |  |
|  | Gleason score of primary tumor prostatectomy (Grade group) | % | (N) |
|  | 3 + 3 (1) | 14.3 | (1) |
|  | 4 + 3 (3) | 28.6 | (2) |
|  | 4 + 5; 5 + 4 (5) | 57.1 | (4) |
|  | Prior systemic therapy * | % | (N) |
|  | Abiraterone | 71.4 | (5) |

|  |  |  |
| --- | --- | --- |
| Enzalutamide | 28.6 | (2) |
| Docetaxel | 42.9 | (3) |
| Sipuleucel-T | 14.3 | (1) |
| Prostvac | 28.6 | (2) |

We performed ULP-WGS on plasma collected prior to radical prostatectomy in the 112 patients with localized disease (Fig. 1a) to an average depth of 0.36× (range: 0.19× to 0.74×). Plasma from the first 40 patients was collected in K_2_-EDTA tubes, while the remainder of blood samples were collected in Streck Cell-Free DNA blood collection tubes (BCTs). With the exception of systemic artifacts in chromosomes 5, 6, 8 and 12 from all plasma collected in the EDTA tubes, no SCNAs were detected. Similarly, in the Streck-collected samples, no SCNAs were detected except for random sequencing artifacts in 5 patients. Because average percent tumor content (PTC) is calculated based on all SCNA events, removal of these artifacts resulted in no calls of PTC. The majority of patients (95 out of 112) had PSA levels ≤ 10 ng/mL. Even the patient with the highest PSA in the entire cohort, 43.63 ng/mL, failed to show non-artifactual SCNAs typical of prostate cancer. Consequently, PTC and percent genome altered (PGA) were indeterminate for the localized cohort.

**Figure 1.**
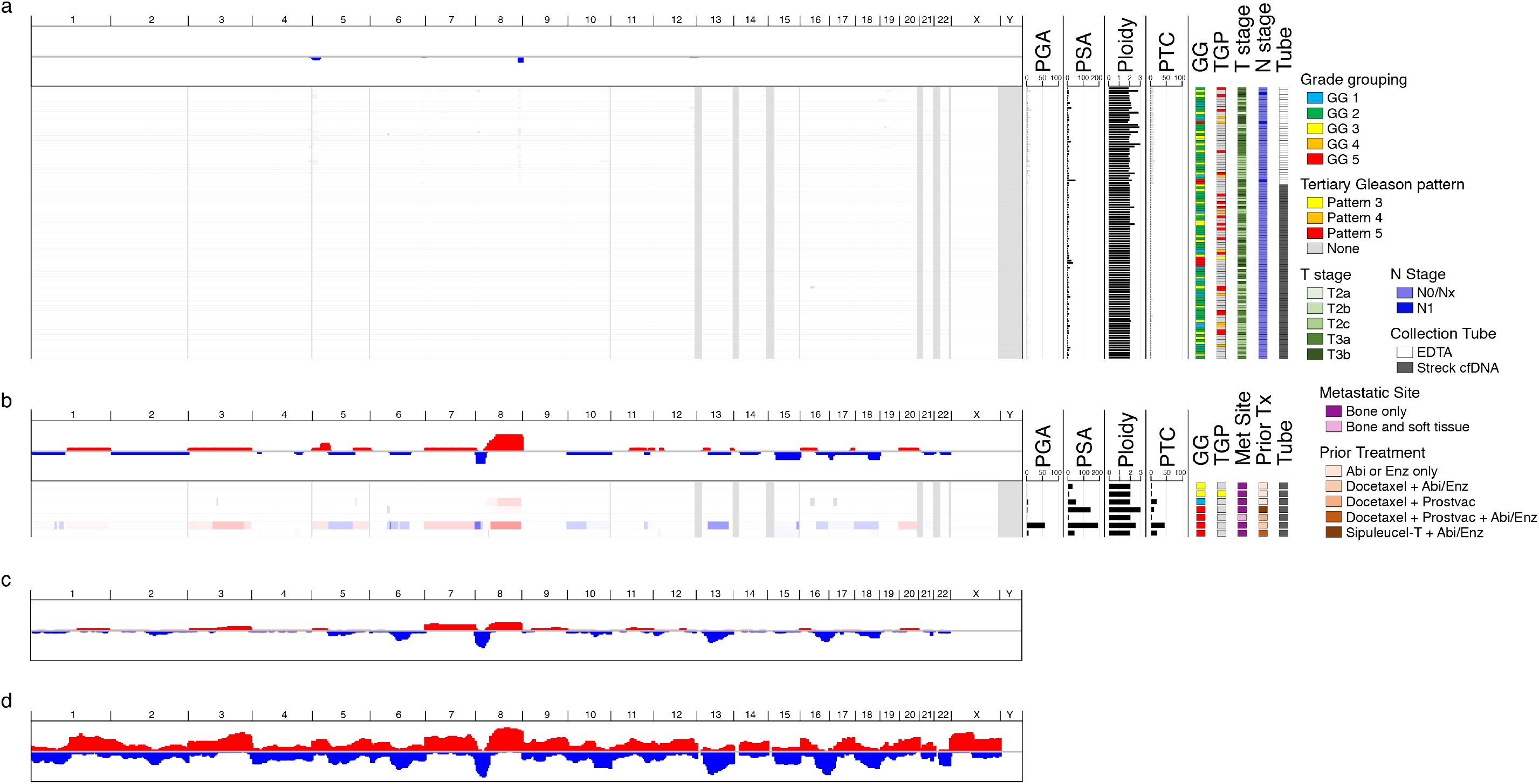
Ultra-low pass whole genome sequencing of circulating tumor DNA. (a) Somatic copy number alteration (SCNA) profile of ctDNA from patients with localized prostate cancer, encompassing National Comprehensive Cancer Network risk groups of low, intermediate-favorable, intermediate-unfavorable, high, and very high-risk disease (*N* = 112). Gray bars represent PGA and PTC values prior to artifact removal. Ploidy values are uncorrected. (b) SCNA profile of ctDNA from patients with radiographically-confirmed metastatic castration-resistant prostate cancer (*N* = 7). PGA: percent genome altered (%); PSA: prostate specific antigen (ng/mL); PTC: percent tumor content (%); GG: International Society of Urological Pathology grade grouping; TGP: tertiary Gleason pattern; Abi: abiraterone acetate plus prednisone; Enz: enzalutamide. (c) SCNA profile of patients in the prostate The Cancer Genome Atlas [24] cohort (*N* = 333). (d) SCNA profile of patients in the Prostate Cancer Foundation-Stand Up to Cancer [44] cohort (*N* = 150).

In contrast, four of the seven patients with metastatic disease had plasma harboring substantial quantities of ctDNA as detected by ULP-WGS (Fig. 1b). The SCNA profile of this cohort was similar to that of The Cancer Genome Atlas (TCGA) prostate cohort (Fig. 1c) and even more similar to the metastatic Prostate Cancer Foundation-Stand Up to Cancer (PCF-SU2C) cohort (Fig 1d). The patient with the highest PTC (30.66%) had a PSA of 190.9 ng/mL, while the patient with the lowest detectable PTC (6.8%) had a PSA of 144.2 ng/mL. In contrast, the lowest metastatic PSA associated with detectable tumor was 42.3 ng/mL, corresponding to 13.94% tumor content. We therefore conclude that ULP-WGS is not sensitive for the detection of SCNAs in the plasma of patients with localized prostate cancer and that PSA in the localized setting is a poor surrogate for likelihood of detecting ctDNA by ULP-WGS.

### Requirements for a patient-specific assay

Development of primary prostate cancer is driven primarily by structural rearrangements and SCNAs; hotspot point mutations in oncogenes and tumor suppressors, such as *HRAS* and *TP53*, are rare [24]. Commercial off-the-shelf ctDNA tests are focused on these recurrent mutations, limiting their utility for detecting ctDNA in primary prostate cancer. Even the most recurrent mutation in primary prostate cancer, at codon 133 of *SPOP*, occurs in fewer than 5% of tumors [24]. Presuming that mutation events that occur early in a tumor’s natural history are present in all daughter cells, truncal passenger mutations would be present in ctDNA and therefore might be used to ctDNA on a per-patient basis.

When mapped, prostate cancers branch substantially at their index lesion (Fig. 2a), such that repeated sampling of multiple tumor regions (Fig. 2b) is needed to empirically infer mutations that are shared by all or most tumor lesions, and thus would be candidates for detection in. Our approach attempts to identify such mutations through several steps, as illustrated in Figure 2c: (1) immunohistochemistry (IHC) was performed on serial sections of multiple blocks of tumor tissue from each patient, (2) laser capture microdissection (LCM) was used to isolate histologically distinct foci from which DNA was extracted; (3) extracted DNA was subjected to WGS and WES; and (4) WGS and WES data was integrated into tumor phylogenies encompassing both SCNAs and point mutations (Fig. 2d). Point mutations comprising the “trunk” or major “branches” of these evolutionary “trees” were selected for incorporation into the patient specific assay.

**Figure 2.**
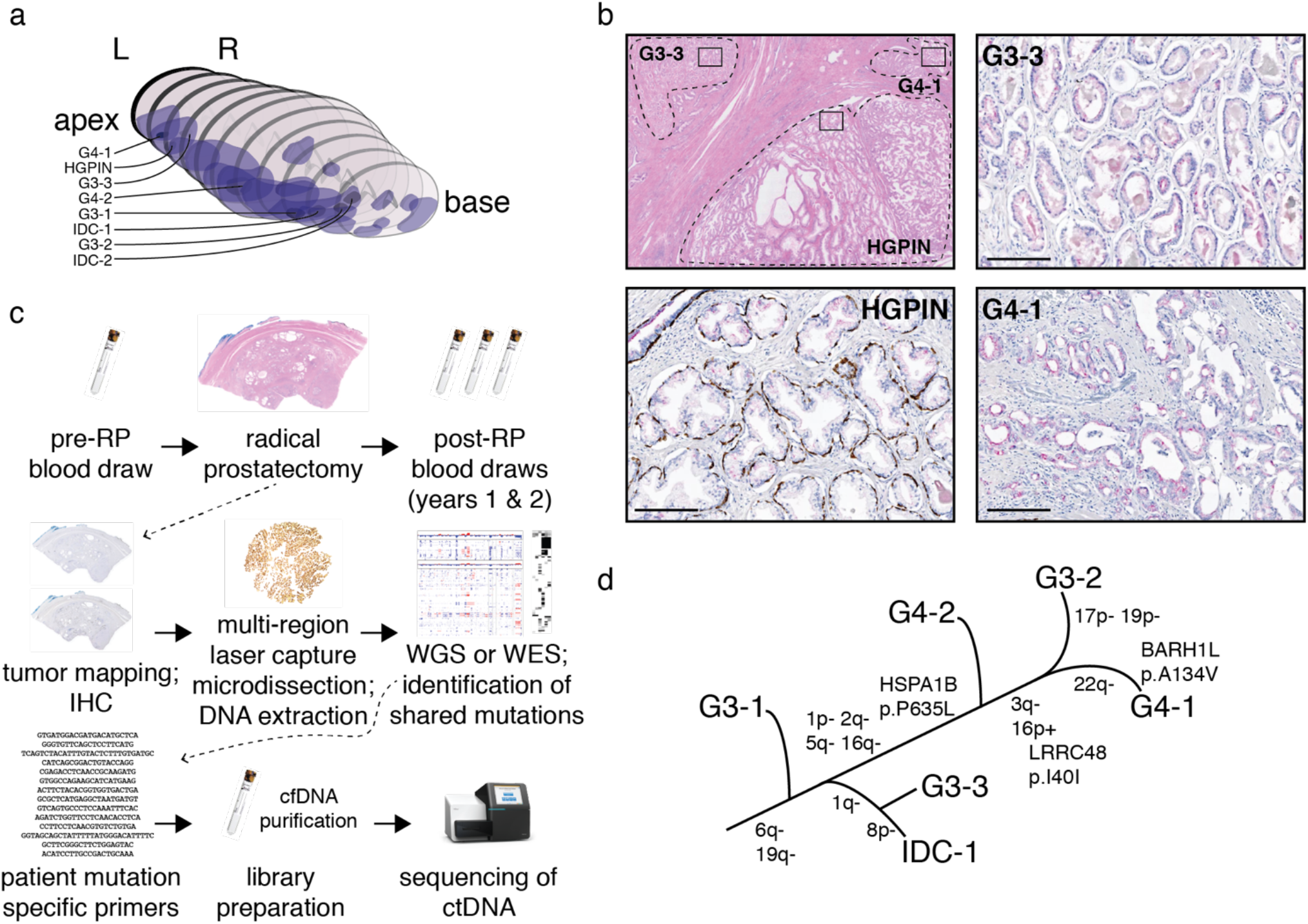
Multi-region sampling of prostate cancer tissue for identification of alleles to be detected in circulating tumor DNA (ctDNA). (a) Representative case demonstrating mapping of tumor throughout the prostate and the selection of distinct histologies that may represent major branches of the tumor system. (b) H&E staining and PIN-4 cocktail immunostaining of three adjacent histologies. Brown chromogen: p63 and cytokeratins 5 and 14; red chromogen: α-methylacyl coenzyme A racemase; bar: 200 μm. (c) General schematic of workflow showing the sequencing of patient tissue for identifying candidate alleles and the retrospective analysis of those alleles in banked plasma from the same patient. (d) Phylogenetic tree of tumor foci from the prostate cancer mapped in (a). While copy number alterations and point mutations are used for establishing the evolutionary tree, only point mutations are sequenced in plasma specimens. RP: radical prostatectomy; IHC: immunohistochemistry; WGS: whole genome sequencing; WES: whole exome sequencing.

### Single molecule detection

Discordance between different commercial tests and even repetitions of standard polymerase chain reaction (PCR) to detect and quantify rare alleles can often be linked to high false positive rates [25, 26]. Consequently, we designed a locus-specific Illumina-compatible library design that incorporated 7-base unique molecular identifiers (UMIs) for tagging individual template molecules. Coupled with analysis scripts that employ heuristics, this design distinguished mutations arising from errors during library preparation from those present in the starting material, making it robust to false positive results. Details of library design and analysis are provided in materials and methods.

To verify and benchmark this design, we first generated target amplicons of 139 basepairs or less (Supp. Table 1), containing eight different heterozygous and homozygous alleles from PC3 and DU145 genomic DNA (gDNA), to serve as synthetic ctDNA. Fresh plasma was obtained from a single male donor with no known cancer diagnoses through the National Institutes of Health Department of Transfusion Medicine. Synthetic ctDNA was spiked into separate 3-mL aliquots of plasma in duplicate for both DU145- and PC3-derived gDNA in approximate copy number amounts spanning six logs (10^1^, 10^2^, 10^3^, 10^4^, 10^5^, and 10^6^) for a total of 24 plasma samples plus four negative controls. Samples were frozen overnight and later thawed for extraction of cfDNA. Two rounds of library preparation were performed per cfDNA sample, for a total of 56 libraries.

As shown in Supplementary Table 2, the assay demonstrated high reproducibility between spike-in targets at similar quantities, with robust detection of mutant alleles at the 10 and 100 spike-in amounts. While the expected limit of detection based on the total possible number of UMI’s was 16,384 (4^7^) template molecules, our observed mean saturation was closer to 1000 molecules (Supplementary Fig. 1), due to the abundance of the wild-type allele which our analysis ignored. Although we spiked in excess quantities of target for the purpose of estimating recovery, yield, and complexity loss during library preparation, this assay demonstrates robust recovery of rare alleles, its intended purpose.

### Positive detection of ctDNA alleles in plasma from patients with metastatic prostate cancer

Tissue biopsies from four out of seven metastatic patients (M03, M04, M06, and M07) with high plasma tumor content, as determined by ULP-WGS (see Fig. 1), were unavailable for sequencing. Whole exome sequencing (WES) was performed on cfDNA from these four cases, employing their matched buffy coat gDNA as a benign control. cfDNA and buffy coat DNA were sequenced to mean on-target depths of 140× and 90×, respectively. As expected, the SCNA profile from exome sequencing generally matched the SCNA profile from ULP-WGS for each patient (Fig. 3a), although the substantially higher resolution of exome sequencing permitted detection of smaller genomic events (Fig. 3b).

**Figure 3.**
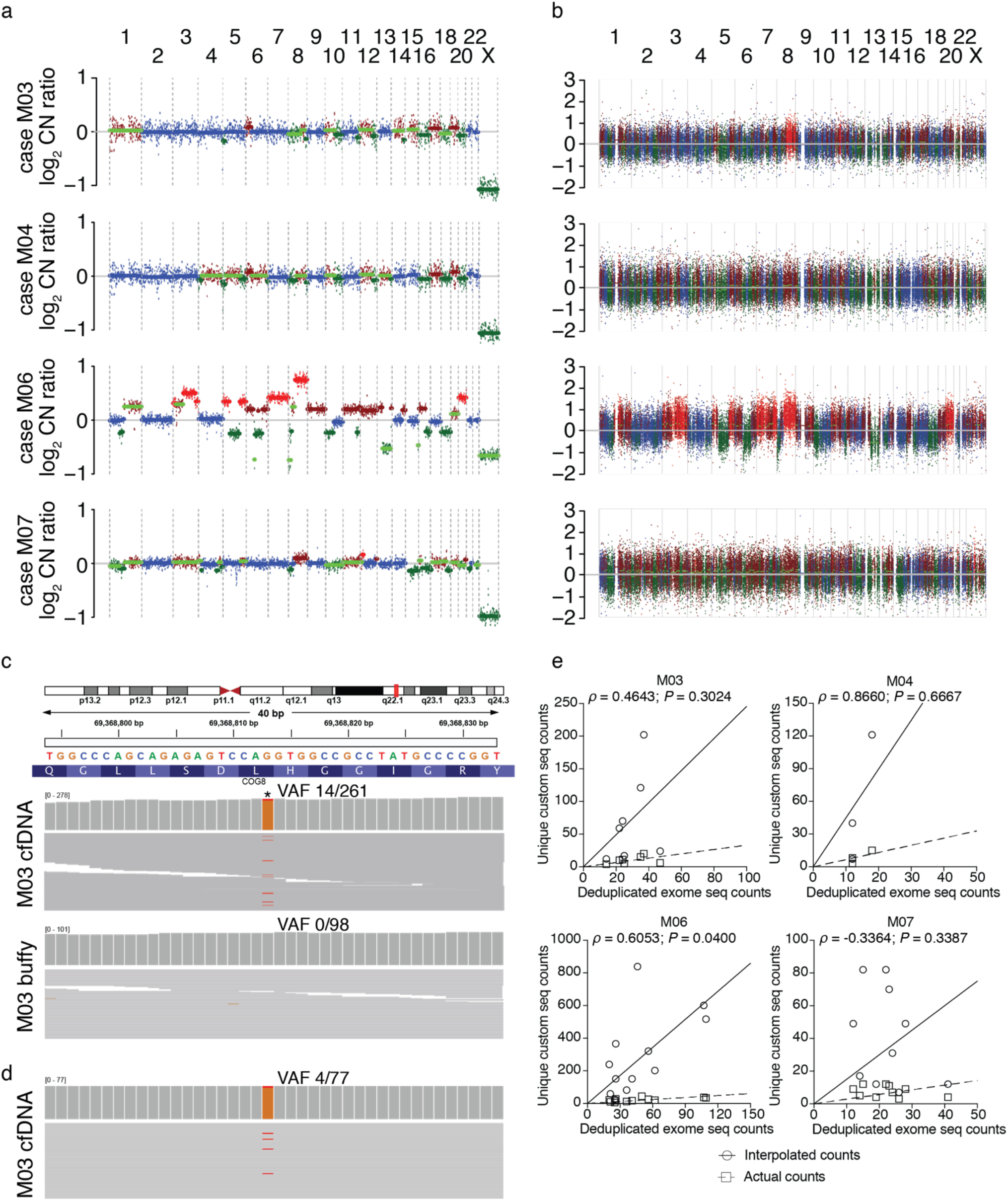
Detection of circulating tumor DNA in plasma from 4 patients with metastatic prostate cancer. (a-b) Plots of log_2_ copy number ratio (cell-free DNA vs buffy coat) for subjects M03, M04, M06 and M07 as determined by ultra-low pass whole genome sequencing (a) and high depth whole exome sequencing (WES) (b). (c) A representative somatic point mutation (g.chr16:69368813G>T) observed in the cfDNA from patient M03 by WES. VAF: variant allele fraction (mutant reads/total reads): * indicates mutant allele. (d) Representative bespoke sequencing detection of mutant g.chr16:69368813G>T allele. (e) Scatter plots showing relationship of actual and adjusted (interpolated) bespoke sequencing mutant read counts versus mutant read counts from exome sequencing. Correlation statistic Spearman’s ρ and *P* values are the same for both actual and interpolated counts).

Importantly, for samples with orthogonally-confirmed ctDNA levels, we generated a catalogue of somatic mutations by comparing each sample with its matched benign control (Suppl. Table 3 and Fig. 3c). Following our protocol for bespoke library design, four sets of primers were generated for the detection of high clonality mutant alleles in each sample (Suppl. Table 4). These primer pairs successfully amplified 36 out of 36 targets from all four cases and detected mutant ctDNA alleles in 32 out of 36 amplified targets (Fig. 3d, Table 2). When back-calculated to an expected molecule number using the spike-in curve fit, raw deduplicated mutant read counts from the patient-specific assay correlated well with the unique mutant read counts from exome sequencing (Fig. 3e).

**Table 2.** Read count data for bespoke cell-free DNA sequencing libraries from four patients with metastatic prostate cancer. Reference and alternate reads count only unique molecules. Read depth: total number of mapped and unmapped reads.

| *Case* | *Mutation* | *Raw locus depth* | *Unique ref alleles* | *Unique alt alleles* | *Library read depth* |
| --- | --- | --- | --- | --- | --- |
| M03 |  |  |  |  | 3417333 |
|  | g.chr5:173473835C>T | 232037 | 101 | 10 |  |
|  | g.chr8:77616607A>T | 272120 | 117 | 11 |  |
|  | g.chr14:23884954_23884955ins | 527471 | 174 | 15 |  |
|  | g.chr16:69368813G>T | 156333 | 73 | 4 |  |
|  | g.chr16:89789061G>C | 208356 | 84 | 5 |  |
|  | g.chr17:7578406C>T | 190463 | 72 | 6 |  |
|  | g.chr19:51885774C>T | 391421 | 128 | 20 |  |
| M04 |  |  |  |  | 2477707 |
|  | g.chr1:153275064T>A | 32293 | 19 | 0 |  |
|  | g.chr1:211527783G>C | 28687 | 16 | 0 |  |
|  | g.chr3:10381924A>G | 416467 | 73 | 15 |  |
|  | g.chr4:190878562G>C | 11247 | 5 | 0 |  |
|  | g.chr8:101721933delT | 15818 | 20 | 0 |  |
|  | g.chr10:87614319A>T | 63914 | 17 | 3 |  |
|  | g.chr19:19375661delC | 266068 | 46 | 8 |  |
| M06 |  |  |  |  | 4365632 |
|  | g.chr2:141607694C>A | 138731 | 57 | 6 |  |
|  | g.chr4:76434490T>A | 233441 | 73 | 17 |  |
|  | g.chr5:112175951_112175952ins | 75879 | 40 | 20 |  |
|  | g.chr7:103801558C>A | 124877 | 54 | 12 |  |
|  | g.chr8:48874964C>A | 213330 | 76 | 28 |  |
|  | g.chr11:104901134C>A | 92355 | 47 | 26 |  |
|  | g.chr12:29642561C>A | 545008 | 128 | 44 |  |
|  | g.chr15:39879635G>C | 91659 | 84 | 17 |  |
|  | g.chr17:38285608C>A | 39445 | 27 | 10 |  |
|  | g.chr19:33288751T>G | 225499 | 93 | 22 |  |
|  | g.chr20:55915197A>C | 134518 | 46 | 37 |  |
|  | g.chr22:29965251G>C | 173584 | 74 | 34 |  |
| M07 |  |  |  |  | 2991208 |
|  | g.chr1:40433489T>A | 23092 | 38 | 4 |  |
|  | g.chr1:153391713G>T | 145392 | 58 | 12 |  |
|  | g.chr1:182811710C>G | 307190 | 98 | 5 |  |
|  | g.chr6:129813480C>A | 254625 | 75 | 11 |  |
|  | g.chr6:130762831T>G | 28261 | 29 | 3 |  |
|  | g.chr7:31918688T>G | 88994 | 67 | 4 |  |
|  | g.chr11:46637383G>T | 88014 | 57 | 9 |  |
|  | g.chr16:55855305G>T | 196877 | 87 | 12 |  |
|  | g.chrX:96328009G>T | 82047 | 38 | 9 |  |
|  | g.chrX:151092687G>T | 20277 | 20 | 7 |  |

Detection of mutant alleles in cfDNA was also positively correlated with raw read count abundance (Suppl. Fig. 2a; Spearman’s *ρ* = 0.5575; *P* = 0.0004). Although interpolation of actual read counts to estimate the number of starting template molecules generally increased the absolute number of alleles reported, the interpolated values were more similar in range to the number of deduplicated exome-seq detected alleles than the actual count (Wilcoxon matched-pairs signed rank test; Suppl. Fig. 2b). We next asked whether lower read count thresholds would impact binary detection (presence/absence) of ctDNA below defined read count thresholds. When sequence reads were downsampled prior to alignment to one million reads per library, or 100,000, 50,000 or 10,000 reads per target, the observed read count was consistently higher than what would be expected due to lower depth (Suppl. Fig. 2b), such that reduction to 10,000 reads per target (>90% downsampling) only reduced mutant allele detection by approximately 50%. At the lowest level of downsampling (10,000 reads per target), all mutant alleles were still detected. Importantly, at sites where mutations were not previously detected, analysis of full sequence output did not reveal mutations that would represent artifacts of library preparation or sequencing. Therefore, from this data we conclude that our patient-specific assay can robustly resequence mutant cfDNA alleles with >93% sensitivity and 100% specificity for the target regions assessed.

### Lack of detection of ctDNA alleles in plasma from patients with localized prostate cancer

With a highly sensitive patient-specific assay robust against false positive results, we applied our ctDNA detection approach to men newly diagnosed with localized disease. Our initial hypothesis was that detection of ctDNA at baseline would predict adverse pathologic features associated with recurrence (such as high Gleason score) or would predict recurrence itself. We selected nine of the 112 localized disease cases, representing a range of Gleason scores, pathologic T-stages, sample ages, baseline PSA levels, and biochemical recurrence statuses (three years or more after prostatectomy), from which to identify clonal markers in plasma (Table 3). A list of microdissected and sequenced foci is given in Supplementary Table 5.

**Table 3.** Clinical and experimental data for the mine patients for tissue sequencing and subsequent circulating tumor DNA assessment from plasma. Gleason score “T”: tertiary pattern; C: Caucasian; AA: African American; WGS: whole genome sequencing; WES: whole exome sequencing; LCM: laser capture microdissected; BCR: biochemically recurrent; R: remission. *Tissue foci from the same block or histologically distinct tumor may have been pooled prior to sequencing. Number does not include additional samples of benign material sequenced as reference controls.

| *Case* | *Gleason score* | *Baseline PSA* | *T-stage* | *Race* | *Patient age (y)* | *BCR status* | *Sample age (y)* | *Sequencing approach* | *Foci LCM* | *Samples sequenced\** |
| --- | --- | --- | --- | --- | --- | --- | --- | --- | --- | --- |
| L001 | 3+4 T5 | 3.38 | T3a | C | 51 | R | 5 | WGS | 5 | 2 |
| L002 | 4+3 | 6.3 | T3a | C | 65 | R | 5 | WGS | 6 | 2 |
| L003 | 3+4 | 8.82 | T3b | C | 66 | R | 5 | WGS | 4 | 2 |
| L004 | 4+3 T5 | 5.8 | T3b | C | 55 | R | 5 | WES | 9 | 9 |
| L015 | 5+5 T4 | 1.6 | T3b | C | 69 | BCR | 4 | WES | 5 | 5 |
| L039 | 5+4 | 43.63 | T3b | C | 68 | R | 4 | WES | 13 | 11 |
| L040 | 4+5 | 6.8 | T2c | AA | 62 | R | 4 | WES | 7 | 7 |
| L107 | 3+3 T4 | 2.8 | T2c | C | 56 | R | 3 | WES | 7 | 4 |
| L108 | 3+4 | 17.3 | T3a | AA | 64 | R | 3 | WES | 15 | 8 |

Prior to LCM, we performed IHC against ERG to select concordantly positive or negative foci (see Suppl. Table 5). We previously established that chromosomal breakpoints serve as a definitive clonal marker in TMPRSS2:ERG fusion positive tumors [27]. In only one (L001) of two ERG-positive cases (L001 and L003) for which WGS was performed did we successfully read through the *TMPRSS2-ERG* breakpoint (Suppl. Fig. 3a-c). However, even a nested PCR approach failed to amplify the fragment of DNA containing the breakpoint from plasma (Suppl. Fig 3d-e).

We therefore employed the approach illustrated in Figure 2, in which we integrated SCNA and mutation clonality data from multiple foci (Suppl. Fig. 4) to identify point mutations as either trunks, branches, or leaves of a given tumor “tree;” truncal mutations were shared by all foci, branch mutations were shared by most or some foci, and leaf mutations were unique to a given focus. The complete list of somatic mutations considered for this analysis is in Supplementary Table 6, and the mutations selected for bespoke sequencing ctDNA analysis are in Supplementary Table 7. Despite high specificity and coverage, no mutated alleles indicative of ctDNA were detected from the cfDNA sampled prior to radical prostatectomy (Table 4). Surprisingly, any ctDNA that may have been present from patient L015, who had Gleason 10 prostate cancer and subsequently recurred, was below the limit of detection for both ULP-WGS (see Fig. 1) and allele-specific measurement (see Table 4 and Suppl. Table 9).

**Table 4.** Summarized read count data for bespoke cell-free DNA sequencing libraries from nine patients with localized prostate cancer, sampled from pre-operative plasma. Reference and alternate reads count only unique molecules. Read depth: total number of mapped and unmapped reads. Data is shown as mean (95% confidence interval). Locus level data is shown in Supplementary Table 9.

| *Case* | *Target #* | *Locus depth* | *Unique ref alleles* | *Unique alt alleles* | *Library read depth* |
| --- | --- | --- | --- | --- | --- |
| L001 | 11 | 49480 (21444 – 77516) | 14 (8 – 20) | 0 (0 – 0) | 954091 |
| L002 | 10 | 12088 (6422 – 17754) | 20 (16 – 25) | 0 (0 – 0) | 259916 |
| L003 | 8 | 28524 (19557 – 37490) | 11 (8 – 14) | 0 (0 – 0) | 421763 |
| L004 | 6 | 67143 (34678 – 99608) | 17 (10 – 23) | 0 (0 – 0) | 790288 |
| L015 | 11 | 37823 (23040 – 52605) | 8 (6 – 10) | 0 (0 – 0) | 732729 |
| L039 | 8 | 64083 (36120 – 92046) | 60 (49 – 71) | 0 (0 – 0) | 828342 |
| L040 | 11 | 36594 (4677 – 68511) | 8 (3 – 13) | 0 (0 – 0) | 733865 |
| L107 | 5 | 100173 (44033 – 156313) | 39 (30 – 49) | 0 (0 – 0) | 1119431 |
| L108 | 13 | 19129 (12114 – 26143) | 83 (67 – 100) | 0 (0 – 0) | 418931 |

Finally, we asked whether we could detect ctDNA in this same group of patients following radical prostatectomy when PSA levels are low prior to biochemical recurrence. Although only one patient in this cohort has recurred to date, ctDNA was not detected in any of the nine patients over multiple time points (Table 5 and Suppl. Table 10). Taken together, we conclude that although allele-specific detection is a robust approach for identifying ctDNA alleles in metastatic prostate cancer patients, it is inferior to the sensitivity of PSA testing in a localized prostate cancer population for measuring disease burden.

**Table 5.** Summarized read count data for bespoke cell-free DNA sequencing libraries from nine patients with localized prostate cancer, sampled from post-radical prostatectomy plasma. Reference and alternate reads count only unique molecules. Read depth: total number of mapped and unmapped reads. Data is shown as mean (95% confidence interval) of the same targets shown in Table 4. PSA was measured concurrently on a separate blood sample at the timepoint shown. Locus level data is shown in Supplementary Table 10.

| *Case* | *Days post-RP* | *PSA (ng/mL)* | *Locus depth* | *Unique ref alleles* | *Unique alt alleles* | *Library read depth* |
| --- | --- | --- | --- | --- | --- | --- |
| L001 | 122 | < 0.1 | 35052 (13613 – 56492) | 20 (11 – 28) | 0 (0 – 0) | 725843 |
|  | 234 | < 0.1 | 42892 (16574 – 69211) | 15 (8 – 22) | 0 (0 – 0) | 851599 |
|  | 353 | < 0.1 | 43098 (14429 – 66366) | 16 (7 – 24) | 0 (0 – 0) | 884849 |
|  | 472 | < 0.03 | 45855 (18682 – 73027) | 24 (13 – 35) | 0 (0 – 0) | 964686 |
| L002 | 106 | < 0.01 | 43847 (23681 – 64013) | 35 (25 – 45) | 0 (0 – 0) | 1007266 |
|  | 246 | < 0.1 | 31569 (15072 – 48065) | 16 (11 – 20) | 0 (0 – 0) | 678782 |
| L003 | 12 | 0.5 | 40493 (28782 – 52204) | 20 (15 – 24) | 0 (0 – 0) | 579156 |
|  | 96 | 0.1 | 57914 (48885 – 66943) | 18 (15 – 20) | 0 (0 – 0) | 821136 |
|  | 418 | < 0.1 | 52945 (38325 – 675660) | 20 (16 – 24) | 0 (0 – 0) | 833626 |
|  | 572 | < 0.1 | 56594 (43599 – 69589) | 23 (20 – 25) | 0 (0 – 0) | 792204 |
|  | 719 | < 0.03 | 72892 (42823 – 102960) | 13 (9 – 17) | 0 (0 – 0) | 995160 |
| L004 | 85 | < 0.1 | 72629 (19149 – 126109) | 18 (8 – 28) | 0 (0 – 0) | 734314 |
|  | 120 | < 0.1 | 96965 (52280 – 141650) | 20 (14 – 25) | 0 (0 – 0) | 1007444 |
|  | 295 | < 0.1 | 85976 (36831 – 135120) | 25 (15 – 36) | 0 (0 – 0) | 895012 |
|  | 598 | < 0.03 | 96241 (53513 – 138969) | 27 (17 – 37) | 0 (0 – 0) | 976471 |
| L015 | 92 | < 0.1 | 74589 (29999 – 119179) | 19 (10 – 28) | 0 (0 – 0) | 1531476 |
|  | 183 | < 0.1 | 30260 (12057 – 48463) | 6 (4 – 8) | 0 (0 – 0) | 593773 |
|  | 239 | < 0.1 | 38473 (18679 – 58267) | 6 (4 – 8) | 0 (0 – 0) | 785887 |
| L039 | 106 | 0.5 | 42633 (22160 – 63107) | 52 (43 – 61) | 0 (0 – 0) | 573305 |
|  | 169 | < 0.1 | 46211 (23761 – 68661) | 47 (40 – 55) | 0 (0 – 0) | 551245 |
|  | 435 | < 0.03 | 43900 (22122 – 65688) | 55 (42 – 67) | 0 (0 – 0) | 528331 |
| L040 | 187 | < 0.1 | 45218 (18290 – 72146) | 12 (7 – 18) | 0 (0 – 0) | 933997 |
|  | 305 | < 0.1 | 40666 (21572 – 59761) | 26 (17 – 35) | 0 (0 – 0) | 926748 |
|  | 396 | < 0.03 | 30376 (5 – 60747) | 5 (1 – 8) | 0 (0 – 0) | 760326 |
| L107 | 128 | < 0.03 | 72126 (41458 – 102794) | 784 (583 – 986) | 0 (0 – 0) | 670825 |
|  | 213 | < 0.03 | 124569 (58162 – 190976) | 75 (60 – 90) | 0 (0 – 0) | 917898 |
|  | 367 | < 0.03 | 92813 (46559 – 139067) | 74 (63 – 85) | 0 (0 – 0) | 743457 |
| L108 | 105 | 0.06 | 46690 (29164 – 64215) | 81 (69 – 92) | 0 (0 – 0) | 1028891 |
|  | 208 | 0.05 | 41151 (25712 – 56589) | 105 (86 – 124) | 0 (0 – 0) | 1024881 |

## DISCUSSION

In light of concerns that PSA levels simply reflect tumor volume, rather than grade, and that they may fail to detect androgen receptor low or indifferent tumors, PSA measurement remains an excellent biomarker for treatment response and it is the gold standard for diagnosing biochemical recurrence after primary therapy [28]. In our study, we hypothesized that higher grade, more poorly-differentiated cancers could be distinguished from indolent tumors based on detection of ctDNA in preoperative plasma. We further hypothesized that plasma from patients with more aggressive tumors that ultimately recurred would also harbor ctDNA that could be detected preoperatively, or postoperatively prior to biochemical recurrence defined by PSA. Using unbiased ULP-WGS, we were unable to detect ctDNA in localized prostate cancer before surgery from patients with a wide range of PSA levels and tumor aggressiveness. Our allele-specific assay, which is sensitive to as few as 10 mutant alleles of spiked-in DNA, similarly did not detect either ctDNA from preoperative plasma or from plasma prior to biochemical recurrence. In contrast, both assays detected ctDNA in patients with metastatic prostate cancer.

If ctDNA levels were directly proportional to PSA levels, then a subset of localized prostate cancer patients with higher PSA would have been expected to have detectable ctDNA [29]. Indeed, in our metastatic cohort, three patients with PSA levels < 30 ng/mL failed to show any SCNAs by ULP-WGS, with the remainder (including one patient with PSA of 51.34 ng/mL) having ctDNA detectable by ULP-WGS, WES, and allele-specific sequencing. However, amongst the localized cohort, even the patient with the highest preoperative PSA (L039; 43.63 ng/mL) failed to harbor detectable ctDNA. This finding suggests that intrinsic differences between primary and metastatic prostate cancer, including the kinetics of ctDNA shedding and turnover, low proliferative rate of localized disease, and poor proximity to vasculature relative to metastases may result in degradation of cfDNA before it reaches circulation.

Because ctDNA potentially represents a pool of multiple subclones shedding cfDNA, alleles detected in ctDNA may only represent the most clonal and truncal of alterations, especially when the percentage of ctDNA in total cfDNA is low [16]. To address this challenge, we reconstructed tumor phylogenies from genome and exome sequencing of tissue in order to select alleles representing the major subclones that would be present at the time of surgery and further mediate relapse. Prior to executing these experiments, we developed and tested a bespoke patient-specific, allele-specific sequencing assay that satisfied requirements for reproducibility, accuracy, sensitivity, and specificity [16, 30, 31]. This assay consistently detected spiked-in alleles and showed 100% concordance to unbiased whole exome sequencing of the same sample at very high coverage. However, after applying this assay to both preoperative and postoperative plasma, we found that the lack of detection of clonal alleles in ctDNA was not predictive of adverse final pathology, recurrence or metastasis.

There is an important limitation of this finding. Although rare, some prostate cancer patients recur with tumors that were only a minor subclone at the time of radical prostatectomy [32]. In our study, we focused on the index lesion as the tumor system most likely to drive relapse. Given the prospective and unselected population of our cohort, the vast majority have remained in remission following surgery, with the only one recurrent tumor (patient L015) having undergone in-depth primary tumor sequencing. Moreover, we were unable to acquire metastatic tissue from this patient to sequence and compare to the targets selected from the prostatectomy. Consequently, it is possible that the clone driving metastasis was independent of the tissue sequenced. Despite using the most sensitive allele-specific assay possible, design of these assays was based on comprehensive tumor sampling. Therefore, we cannot state with absolute certainty that allele-specific analysis assessed the correct clone and that ctDNA levels in patients were below limits of detection.

There have been a limited number of published studies that evaluate cfDNA as a biomarker prognostic of advanced disease in the localized prostate cancer setting [33–35]. The largest of these studies to date examined the total burden of cfDNA and ctDNA by hypermethylation of the *GSTP1* promoter in DNA extracted from the serum of 192 patients [34]. Although *GSTP1* hypermethylation in serum cfDNA was increased in the recurrent and metastatic populations compared to indolent prostate cancer, contribution of *GSTP1* equivalents in serum from normal tissue affected by oligometastases may have contributed to this finding because *GSTP1* hypermethylation is not a tumor-specific event [34]. Moreover, the PCR assay used for detecting circulating *GSTP1* amplifies a DNA fragment in far excess of the ~165 bp ctDNA fragment, suggesting it is of nontumor origin despite reflecting increased tumor aggressiveness [34].

Bespoke approaches to detect ctDNA from urothelial and colorectal cancers have demonstrated success in risk stratification and therapy monitoring. In a cohort of 68 patients with muscle invasive bladder cancer, a personalized assay to sequence somatic variants as markers of ctDNA in preoperative plasma was highly prognostic for recurrence following cystectomy [36]. Amongst recurrent patients, ctDNA detected prior to chemotherapy also tracked with worse overall survival [36]. Similar successes were achieved in a cohort of 130 colorectal cancer patients, in which a personalized ctDNA detection assay detected ctDNA in 88.5% of preoperative plasma, and 70% of patients with detectable ctDNA at the start of adjuvant chemotherapy subsequently recurred [21]. The striking difference between our findings and these reports from bladder and colorectal cancer cohorts may reflect some of the same differences between primary and metastatic prostate cancer with respect to ctDNA shedding, including cell proliferation rate and proximity to vasculature.

Nonetheless, to the best of our knowledge, this is the first comprehensive analysis to conclude definitively that somatic mutation and copy number alterations in cfDNA do not effectively measure ctDNA levels in an untreated localized prostate cancer cohort. Because these locus-level analyses of individual genomes are below the limits of detection, other circulating nucleic acid analytes may be more representative of phenotype and thus offer better detection characteristics. Circulating tumor cells, circulating cell-free miRNA, circular RNA, posttranscriptionally modified RNA species, and genome-wide tissue-of-origin patterns of DNA methylation do not correlate 1:1 with tumor cell number, and thus may give a much greater signal than allele-dependent assays for the early noninvasive detection of aggressive and potentially recurrent prostate cancer [36, 37].

## MATERIALS AND METHODS

### Study approval

This research was conducted in accordance with the principles of the Declaration of Helsinki. The collection and analysis of plasma, tissue and demographic data from patients with localized prostate cancer was approved by the Beth Israel Deaconess Medical Center and Dana-Farber/Harvard Cancer Center Institutional Review Boards, under protocol numbers 2010-P-000254/01 (BIDMC), 15-008 (DFHCC), and 15-492 (DFHCC). The collection of plasma and demographic data from patients with metastatic prostate cancer was approved by the NIH Institutional Review Board, under protocol number 02-c-0179. All patients provided informed consent prior to participating in tissue procurement protocols.

### Blood collection

Blood was obtained from patients diagnosed with localized prostate cancer pre-or perioperatively to radical prostatectomy, and again at intervals coinciding with urology follow-up visits. At each time point, 8-10 mL of whole blood was collected into K_2_-EDTA Vacutainer (Becton Dickinson) or Cell-Free DNA BCTs (Streck) following recommended guidelines for venipuncture, draw order, and inversion. K_2_-EDTA-collected blood was stored on ice and processed within one hour of collection. Cell-Free DNA BCT-collected blood was stored at room temperature and processed within three days of collection (if shipped from DFHCC to NIH) or within 8 hours of collection (if processed in DFHCC).

Blood was obtained from patients with metastatic castration-resistant prostate cancer following progression on therapy. At a single time point prior to receiving subsequent therapy, 8-10 mL of whole blood was collected into Cell-Free DNA BCTs as described above. Sample processing occurred within eight hours of collection.

### Blood processing

Plasma separation followed a two-spin protocol. Initial centrifugation was performed on an Eppendorf 5430R centrifuge with an F-35-6-30 rotor, for 20 minutes at 300 × *g* at 22 °C with ramping speed set to low. Following initial separation, the upper plasma layer was transferred to a new tube, leaving a few mm of plasma above the buffy coat interface. The buffy coat was then transferred to a new tube and stored at −80 °C. The plasma was centrifuged again, at 5,000 × *g* at 22 °C for 10 minutes, with ramping speed set to high. The plasma supernatant resulting from this centrifugation was aliquoted into new tubes and stored at −80 °C.

### DNA extraction from blood samples

gDNA was extracted from 10-100 μL of buffy coat using the DNeasy Blood and Tissue Kit (Qiagen) following the manufacturer’s protocol with two elutions of buffer AE (100 μL per elution). Quality control of buffy coat DNA was performed by assessing double-stranded DNA concentration using PicoGreen (Life Technologies). cfDNA was extracted from plasma using the QIAamp Circulating Nucleic Acid Kit (Qiagen) following the manufacturer’s protocol with minor modifications: buffer ACB was incubated with lysate for 10 minutes on ice and elution was performed twice with buffer AVE (25 μL per elution) following a five minute incubation at room temperature. Quality control of cfDNA was performed by confirming the presence of single or double nucleosome peaks on TapeStation (Agilent) with D1000 ScreenTape, corresponding to ~165 and ~330 bp representing mononucleosome and dinucleosome DNA equivalents, respectively.

### Tissue histology

Radical prostatectomy specimens were grossed, formalin-fixed and paraffin embedded (FFPE) according to standard procedures. Hematoxylin & eosin (H&E) stained slides were reviewed by a board-certified surgical pathologist following the 2014 International Society of Urological Pathology (ISUP) guidelines. For each case, maps were created to identify the distribution of cancerous regions throughout the entire resected specimen.

### Immunohistochemistry (IHC)

Serial sections of FFPE tissues containing benign or abnormal tissue (as identified during mapping) were cut at 5 μm thickness onto Superfrost Plus (Fisher Scientific). Slides were stained with anti-ERG (Abcam, Cat# ab92513, 1:500 into SignalStain diluent), anti-PTEN (Cell Signaling, Cat#9188L, 1:100 into SignalStain diluent), and PIN-4 cocktail (Biocare Medical, Cat# PPM225DS, ready-to-use) antibodies. SignalStain diluent was from Cell Signaling. Slides were baked for 15 minutes at 60 °C, except for PIN-4 stained slides which were baked overnight at 45 °C, deparaffinized through xylenes, and rehydrated through graded alcohols. Antigen retrieval was performed using a NxGen Decloaker (Biocare Medical), for 15 minutes at 110 °C in Tris-EDTA (Abcam, Cat# ab93684) for PTEN, and for 15 minutes at 110 °C in Diva Decloaker (Biocare Medical, Cat# DV2004MX) for ERG and PTEN. Sections were blocked with hydrogen peroxide (Sigma-Aldrich, Cat# 216763) for five minutes, blocked with Background Sniper (Biocare Medical, Cat# BS966) for 10 minutes for PIN-4 or VectaStain Elite ABC HRP kit (Vector Laboratories, Cat# PK-6101) for PTEN and ERG, and incubated with primary antibody overnight at 4 °C (ERG and PTEN) or for one hour (PIN-4). Secondary labeling was performed using Mach 2 Double Stain for PIN-4 (Biocare Medical, Cat# MRCT525) or the VectaStain Elite ABC HRP kit for ERG and PTEN for 30 minutes. Avidin-biotin complexing was then performed for 30 minutes for ERG and PTEN. Colorimetric detection was achieved using DAB Peroxidase HRP (Vector Laboratories, Cat# SK4100) for ERG and PTEN, Betazoid DAB (Biocare Medical, Cat# BDB2004) for PIN-4, and Vulcan Red Fast Chromogen (Biocare Medical, Cat# FR805) for PIN-4. Counterstaining was performed using Mayer’s Hematoxylin Solution (Sigma Aldrich, Cat# MHS16). PIN-4 stained slides were air-dried. ERG and PTEN stained slides were dehydrated through graded alcohol and cleared in xylenes. Slides were mounted using Permount (Thermo Fisher).

For PTEN, normal glands acted as the positive control, and previously stained slides harboring genomically defined PTEN deficiency status served as negative controls. For ERG, endogenous expression of ERG by endothelial cells acted as the positive control and normal stroma was the negative control. H&E- and IHC-stained slides were scanned with an AxioScan.Z1 (Zeiss) at 20 × magnification (Plan-Apochromat, NA 0.8) with brightfield illumination with slide loaders to accommodate 25 mm × 75 mm slides.

### Laser capture microdissection

Serial sections of tumor tissue (and benign regions uninvolved with tumor) were cut onto metal frame PEN-membrane slides (MicroDissect GmbH), stained with Paradise Stain (Thermo Fisher), and laser capture microdissected using an ArcturusXT Ti microscope (Thermo Fisher) onto CapSure Macro LCM Caps (Thermo Fisher). Slides were stained with H&E, ERG, PTEN, and PIN-4, scanned and visualized using ZEN Browser (Zeiss) on an adjacent monitor as references. Per focus, 50,000 to 100,000 cells were captured. For each cap, a photomicrograph was acquired for the purposes of estimating tumor cell purity in each sample.

### DNA extraction from tissue

Microdissected cells adhered to caps were lysed and DNA purified using the QIAamp DNA FFPE Tissue Kit (Qiagen), according to the manufacturer’s instructions, as previously described [38]. DNA yields were quantified using PicoGreen reagent.

### WGS and WES of tissue DNA

gDNA was sheared using acoustic sonication (Covaris). For WGS target genomic fragment sizes were >300 bp and selected by Pippin Prep (Sage). Following end-repair and A-tailing, modified Illumina adaptors containing 7-base inline UMIs at the 3′ end of each adaptor were ligated to each library insert. Libraries were sequenced on a HiSeq X10 (Illumina) instrument to a target depth of 30-40× coverage.

For WES, target fragments were sonicated to a target size of 200 bp and were selected by AMPure XP SPRI beads (Beckman Coulter). WES libraries were prepared using the SeqCap EZ Exome Kit v3 (NimbleGen) or the SureSelect Human All Exon V7 Low Input Exome kit (Agilent). Equimolar pooled libraries were sequenced on HiSeq 2000 and 4000 instruments (Illumina) to a targeted on-bait depth of 150×. Agilent libraries included an additional R3 read to sequence the 10-base UMI.

### Ultra-low pass (ULP-) WGS of plasma DNA

10-100 ng of cfDNA from plasma samples (corresponding to approximately 10 μL of eluate) or 100 ng of gDNA was assembled into paired-end libraries using the NEBNext Ultra II DNA Library Prep Kit (New England BioLabs). Approximately 60 libraries were pooled per lane prior to sequencing on a HiSeq 4000 machine to a target depth of 0.5×.

### WES of plasma DNA

10-100 ng of cfDNA from plasma samples (corresponding to approximately 10 μL of eluate) or 100 ng of gDNA was assembled into exome sequencing libraries using the SureSelect Human All Exon V7 Exome kit (Agilent). DNA samples were pooled to achieve an on-bait depth of 100× for the buffy coat gDNA and 300× for the plasma cfDNA.

### Analysis of WGS and WES

For ULP-WGS, pass-filter FASTQ files were aligned to the b37-decoy version of the human genome using BWA-MEM version 0.7.17, sorted, duplicate-removed using PICARD, and base-recalibrated using GATK version 4.0.5.2. Whole genome copy number profiles and tumor content estimates were generated using the ichorCNA package [19] in R/Bioconductor with one megabase resolution.

For WGS from tissue specimens, pass-filter FASTQ files were parsed to identify the 7-base UMI at the beginning of each R1 and R2 read, and the UMIs were stored in separate files with their read names. UMI-removed FASTQ files were aligned to b37-decoy using BWA-MEM version 0.7.10 and the UMI sequences were appended to the read names for all primary and secondary alignments. PCR duplicates arising from repetitive UMIs with the same mapping were identified using UMITools version 2.1.1 and the corresponding read flags were updated with SAMTOOLS version 1.2 to reflect duplicate marking.

For WES from tissue specimens, pass-filter FASTQ files were adaptor-trimmed using Trimmomatic version 0.27 [39] (for NimbleGen libraries) or SureCall Trimmer version 4.0.1 (Agilent, for Agilent libraries). Trimmed FASTQ files were aligned to b37-decoy using BWA-MEM 0.7.17 and sorted using PICARD. Duplicate marking was performed using PICARD for NimbleGen and LocatIt version 4.0.1 (Agilent) for Agilent libraries, respectively. For WES from blood specimens, pass-filter FASTQ files were adaptor-trimmed using Trimmomatic version 0.27. Trimmed FASTQ files were aligned to b37-decoy using BWA-MEM 0.7.17 and sorted and duplicate marked using PICARD. Following duplicate marking, WES and WGS BAM files were quality-score recalibrated using GATK version 4.0.5.2. For all samples, alignment and duplicate marking was performed first at the lane level. Then for samples for which libraries were run over multiple lanes, quality-score recalibrated BAM files from multiple lanes were merged and duplicate marking was performed a second time.

Point mutations and small indels were identified using MuTect2 in GATK version 4.0.12.0, first by generating a panel of normals for each sequencing and library design methodology, and then making somatic mutation calls reflecting the read depth at each position (GetPileupSummaries) and cross sample contamination (CalculateContamination), with additional filtering for sequencing artifacts (CollectSequencingArtifactMetrics) and strand bias (FilterByOrientationBias). Somatic mutations were annotated using Oncotator with the April 5, 2016 database and were manually inspected using the realigned output BAM files from MuTect2 in IGV.

Large deletions and chromosomal events (including *TMPRSS2-ERG* fusions) were called from WGS BAM files using Delly version 0.5.9 [40]. Whole genome SCNAs were assessed from WES and WGS BAM files using GATK version 4.0.12.0, with panels of normals for each library design methodology. All BAM files were processed through CollectReadCounts, and the panel of normals was used to smooth read counts for each tumor sample using DenoiseReadCounts and CollectAllelicCounts. ModelSegments was used to segment read depth variably across the entire genome (for WGS) or solely in areas corresponding to hybrid capture (for WES).

PlotModeledSegments was used to generate SEG files for inspection and visualization in IGV and corresponding log_2_ copy number ratios for determining ploidy. The ichorCNA component of the TitanCNA pipeline [41] was used to generate per-sample probe level copy number visualization.

### Selection of mutations for bespoke sequencing

For each point mutation, assessment of clonality was performed both by using CLONET [42] and by calculating the cancer cell fraction (CCF) [12], which represents the percentage of tumor cells in the sequenced sample that harbor the mutation. Mutations clonal to the entire tumor had high clonality and CCF values and were shared by most, if not all, of the tumor foci sequenced. Mutations subclonal to the entire tumor but clonal within each focus also had high clonality or CCF values but were exclusive to the focus of tumor sequenced. For each copy number alteration, assessment of clonality was inferred by examining tumor purity and the calculated log_2_ ratio of each segment. For example, a clonal single-copy deletion would show a purity-adjusted log_2_ ratio of 0.5, so deletions with values between 0.5 and 1 were less clonal. A sliding window of 100 kb was allowed for a shared copy number event between microdissected foci. Focal events were evaluated and manually curated within the context of any larger event.

A simple matrix was constructed that listed each SCNA or point mutation and the focus harboring it. Phylogenetic trees were assembled using the most logical path to incorporate the greatest number of shared events between foci with the highest CCF or clonality scores. In cases of a tie between assigning a subclonal event to two or more foci that diverged, the event with the highest CCF or clonality value was given priority with the assumption that spatial proximity between clones and cross-clone contamination accounted for intrafocal genomic discordance. From these trees, we identified point mutations or indels that comprised the trunk or major branches of the tumor and those which comprised the focus-specific leaves on each branch. As the vast majority of mutations were of unknown significance and corresponded to genes not associated with cancer, preference was given to small indels that were unlikely to arise from sequencing artifacts, as well as non-C/T and A/G point mutations, which are common artifacts of PCR [43].

### Design of locus-specific primers for bespoke sequencing

Genomic coordinates for each point mutation, as well the flanking bases 90 nucleotides 5′ and 3′ were identified. Using parameters to limit the amplicon size to 139 nucleotides, Primer3 (http://primer3.ut.ee/), NCBI PrimerBlast (https://www.ncbi.nlm.nih.gov/tools/primer-blast/) and Ion AmpliSeq Designer (https://www.ampliseq.com) websites gave similar results in locus-specific primer design. All multiplex primer panel designs were cross-checked using *In-Silico* PCR (https://genome.ucsc.edu/cgi-bin/hgPcr) to prevent production of unintended amplicons for each patient-specific batch of targets. Each locus-specific primer pair was similarly assessed for on-target amplification.

To the “forward” locus-specific sequence, 40 bases (5′-TCGTCGGCAGCGTCAGATGTGTATAAGAGACAGNNNNNNN-3′) were appended to the 5′ end consisting of a 33-base adaptor and 7-base degenerate UMI to create tripartite “forward” primer. To the “reverse” locus-specific sequence, a 34-base adaptor (5′-GTCTCGTGGGCTCGGAGATGTGTATAAGAGACAG-3′) was appended to the 5′ end to create a bipartite “reverse” primer.

For each target, locus-specific primers not containing the longer adaptor sequences were also designed. Both types of primers were ordered from Integrated DNA Technologies (IDT). Quality control for primer target amplification and performance in multiplex PCR was assessed by screening each primer pair in singleplex reactions using gDNA from PC3 or DU145 cells (10 ng) as template (see below) using the HiFi HotStart Uracil+ ReadyMix PCR Kit (Kapa Biosystems). Reactions were supplemented with uracil N-glycosylase (0.2 U) and dUTP (290 nM). Reactions were incubated at 37 °C for two minutes, 95 °C for five minutes, and then 35 cycles of PCR (95 °C-55 °C-72 °C for 30 seconds each with seven minutes of final extension at 72 °C). PCR products were mixed with gel loading dye and visualized on ethidium bromide stained 1.5% TAE-agarose gels under ultraviolet transillumination.

Sequencing adaptors (Suppl. Table 11) were modified to contain longer regions of complementarity to bipartite and tripartite oligos and were ordered with NGSO-4 purity from Sigma.

### Control DNA

PC3 and DU145 cells were purchased from the American Type Culture Collection (ATCC); no more than 10 passages of each cell line were used before thawing earlier generation stocks. gDNA from PC3 and DU145 cells were purified using the QIAamp DNA Mini Kit (Qiagen) and sonicated to ~170 bp using a Covaris S2 sonicator to create a synthetic DNA template with similar physical properties to cfDNA.

To generate spike-ins, control DNA (10 ng) was amplified with locus-specific primers in single-plex reactions to enrich for regions of known mutation (see Suppl. Table 1). Individual PCR products were silica column-purified (Qiagen), inspected by TapeStation for their predicted amplicon size, diluted to estimated amounts, and pooled.

To test individual primer pairs for specificity, locus-specific primers were tested in singleplex reactions as described above. Primer pairs not demonstrating specificity for PCR were excluded from a subsequent multiplex reaction. Sheared DNA (50 ng) was used for library construction (see below) to determine the optimum number of cycles for the final reaction, on a per library basis, before using cfDNA from plasma.

### Bespoke sequencing library construction

AirClean 600 (Air Clean Systems) PCR hood workstations were used for all reaction setups. Tripartite forward primers and bipartite reverse primers were resuspended in 1 × TE buffer (pH 8.0, 100 μM). A designated pre-PCR workstation was use for the initial UMI tagging step.

For the UMI tagging reaction for each patient, a mixture of all forward primers was prepared such that each primer was represented at 2 μM concentration. An aliquot of the primer mix (2.5 μL) was combined with uracil N-glycosylase (0.3 U, 2 μL), cfDNA (8 μL), 2 × KAPA HiFi HotStart Uracil+ ReadyMix (15 μL), and dUTP (3.5 μL, 2.5 μM). Samples were incubated in a thermal cycler at 37 °C for two minutes, 98 °C for five minutes, 65 °C for two minutes, and 72 °C for seven minutes. Products from the tagging reaction were cleaned using AMPure XP SPRI beads (54 μL for each 30 μL tagging reaction), following the manufacturer’s recommended protocol for mixing, magnetic separation, ethanol washes, and drying. Elution was performed with deionized nuclease-free water (10 μL) to recover the purified tagged product.

Purified tagged products (8 μL) were combined with ddATP (2 μL, 0.2 mM), 10 × CoCl_2_ (2 μL), 10 × terminal deoxynucleotidyl transferase (Tdt) buffer (2 μL), low TE buffer (4.4 μL), and Tdt enzyme (1.6 μL of 20 U) for a total volume of 20 μL. Samples were incubated for 90 minutes at 37 °C and 15 minutes at 75 °C. Samples were held at 4 °C until clean-up, which was performed using the Select-A-Size Clean & Concentrator Kit (Zymo) according to the manufacturer’s instructions for a 100 bp one-sided cutoff with the following modifications: the last wash step was centrifuged at maximum speed for one minute, and elution was performed with 10 μL Zymo elution buffer with a four-minute incubation. 8 μL of purified product were recovered.

For the amplification of targets using the reverse primers, a mixture of bipartite oligonucleotides was created with 1 μL of each 100 μM primer in a total volume of 100 μL TE, for a final concentration of 1 μM each primer. To 1.25 μL of this primer mix, 3.5 μL of 2.5 μM dUTP, 1 μL of TE, 8 μL of product from the terminal transferase reaction, 1.25 μL of a 2 μM stock of i5 sequencing adaptor (see Suppl. Table 11), and 15 μL of 2 × KAPA HiFi HotStart Uracil+ ReadyMix were added. Samples were incubated in a thermal cycler at 98 °C for five minutes initially and followed by 20 cycles of PCR (98 °C-65 °C-72 °C for 30 seconds each with seven minutes of final extension at 72 °C). Clean-up was performed using the Select-A-Size Clean & Concentrator Kit using the guidelines for a 200 bp one-sided cutoff according to the manufacturer’s protocol with the following modifications: the last wash step was centrifuged at maximum speed for one minute, and elution was performed with 10 μL Zymo elution buffer following a five-minute incubation. 8 μL of purified product were recovered.

Final construction of each sequencing library was performed by additional amplification using both the i5 and i7 sequencing adaptors (see Suppl. Table 11) to prime PCR. All 8 μL of product from the previous step was mixed with 4 μL of 4 μM i7 adaptor stock, 4 μL of 2 μM i5 adaptor stock, 4 μL of TE, and 20 μL of 2 × KAPA HiFi HotStart Uracil+ ReadyMix. Samples were incubated in a thermal cycler at 98 °C for five minutes and followed by 10-30 cycles of PCR (98 °C-72 °C for 30 seconds each with seven minutes of final extension at 72 °C). The exact number of cycles of PCR to use were determined empirically by substituting cell line genomic DNA in the first step of the library preparation protocol, substituting 4 μL of 6 × SYBR green in the final PCR step above, and performing qPCR on a QuantStudio 3 (Thermo Fisher). The number of cycles was determined by cycle threshold value corresponding to the ½-maximum of amplification saturation on a linear curve. Clean-up was performed using the Select-A-Size Clean & Concentrator Kit with a 200 bp one-sided cutoff as described above, but with a 20 μL elution.

Quality control was performed on completed libraries using D1000 ScreenTapes on a TapeStation looking for peak sizes between 250 bp and 350 bp with primer dimer comprising less than 10% of the library. Libraries were then individually quantified for functional concentration using the KAPA Illumina Quantification Kit for NGS, making 1:50,000 dilutions of each library per the manufacturer’s protocol. Libraries were pooled to final sequencing concentrations of 2-10 μM with the goal of achieving a balance of reads across each target, weighing the proportion of each library in the final pool by the number of targets in each library.

### Patient-specific ctDNA library sequencing and analysis

Libraries from plasma cfDNA were sequenced on a MiSeq (Illumina) instrument with the v2 Reagent Kit (500-cycle). Depending on the number of patient samples per pool, 10-30% Phi-X was added to the final library pool. Sequencing was performed with 195 cycles paired-end and dual 8-cycle indexing reads on the P5 and P7 adaptors. Onboard adaptor trimming was turned off. This sequencing strategy intentionally sequenced through the entire insert molecule and into the adaptor on the other side, reading the i5 and i7 indexing adaptor an additional time. Libraries from the spike-in experiment were sequenced on a HiSeq 4000 (Illumina) with 250 cycles paired-end and dual indexing.

Individually barcoded libraries were recovered and filtered for quality using bcl2fastq (Illumina). Pass-filter FASTQ files were trimmed with Trimmomatic version 0.36, preserving the trimmed-off sequences and parsing them for the expected i5 and i7 adaptor based on bcl2fastq binning. Comparing each library to the expected adaptor pair, read pairs corresponding to the incorrect adaptors were discarded. Comparing the forward and reverse trimmed reads (corresponding only to the insert), read pairs showing discordance (i.e. the exact sequence was not the same on the R1 and R2 reads) were discarded. For concordant reads, the R2 was discarded and R1 was retained.

The R1 reads were parsed for the 7-base UMI that was at the 5′ of the R1 read and stored in a separate file with the corresponding read name. The UMI-removed FASTQ was aligned to b37-decoy using BWA-MEM version 0.7.10. UMIs were then reassigned to aligned reads in a matrix and the frequency distribution of each read and its UMI were determined. For example, a given sequence might correspond to 20 different UMIs, with three UMIs representing 5000 reads each, and the remaining 17 UMIs representing 1-10 reads each. UMIs representing less than 10% of a given sequence were more likely to have arisen by PCR or sequencing error than have existed as a degenerate barcode at the time of initial library preparation and UMI tagging. Similarly, depending on the total number of read-UMI families passing this threshold, the number of different sequences for a single UMI could range between 50% and 85% but could not be less than 50% of a single UMI’s paired sequence unless additional error was introduced during PCR. Based on these cutoffs, truly unique read-UMI pairs were identified and the remainder of the R1 reads were marked as duplicates. Unique and duplicate-marked entries were merged back into the SAM file and converted to sorted BAM with PICARD.

BAM files were locally realigned and base score recalibrated using GATK 3.6.0. Unique base calls were identified by running pysamstats over the defined intervals for each target expected in each sample. Cross-sample contamination during library preparation was ruled out by running pysamstats over a different patient’s set of targets that were included during library preparation but would be expected to be absent.

For the spike-in assay, duplicate-marked read counts for 8 distinct targets were averaged across two independent experiments performed in duplicate with two different pools of synthetic PC3 or DU145 template at six different concentrations of spike-in. Data were fit to a nonlinear hyperbolic curve (see Suppl. Fig. 1) for the purpose of creating a standard curve for interpreting read counts from experimental data and identifying where allele counts become saturating. For duplicate-marked absolute mutant allele counts from cfDNA, actual starting molecule numbers were interpolated using the curve fit equation.

### Statistics

Statistical analyses were performed using GraphPad Prism 8 for Mac. Statistical tests used and relevant variables are indicated in the legend of each figure.

## ACKNOWLEDGMENTS

The authors gratefully acknowledge the patients and the families of patients who contributed to this study. DNA sequencing was performed at the Center for Cancer Research (CCR) Genomics Core, the CCR Illumina Sequencing Facility, and the CCR Genomics Technology Laboratory. The authors acknowledge technical assistance from Cesar Vazquez, Stephan Duggan, Sean Gerrin, and Carla Calagua. Portions of this research utilized the computational resources of the NIH HPC Biowulf cluster.

## IN MEMORIAM

This work is dedicated to the memory of Dr. Valery Bliskovsky, who made invaluable contributions to this project.

## AUTHOR CONTRIBUTIONS

Design and conduct of the clinical study: P.C., G.J.B., S.P.B., H.Y., A.G.S.; Tissue collection and histology: R.L., P.C., H.Y., N.C.W., S.W.; Blood collection and plasma isolation: S.T.H., N.T.T., S.Y.T., O.S.V., R.J.S., N.V.C., A.R.S., F.K., J.L.G., G.J.B., S.P.B.; Immunohistochemistry and microdissection: S.T.H., S.Y.T., R.J.S., N.V.C., R.A.; Data collection: D.J.E., R.J.S., G.J.B., H.Y.; Library construction and NGS: S.T.H., S.Y.T., N.T.T., S.S., A.R.S.; Data analysis: S.T.H., S.Y.T., N.T.T., A.G.S.; Manuscript preparation: S.T.H., A.G.S.

## DATA AVAILABILITY

Sequence data has been deposited into dbGaP, accession ID phs001813.v1.p1.

## Notes

Financial support: This work was supported by NIH grants (DH/HCC SPORE P50 CA090381 to A.G.S. and S.P.B.), the Prostate Cancer Foundation (Young Investigator Awards to S.W., F.K., H.Y. and A.G.S.), the Department of Defense Prostate Cancer Research Program (W81XWH-15-1-0136 and W81XWH-15-1-0710 to A.G.S.; W81XWH-17-1-0350 to D.J.E.) and the Intramural Research Program of the NIH, National Cancer Institute.

Conflicts of interest: The authors have no conflicts of interest to declare.

#### Summary of Updates

Figures were omitted from the initial upload.

