## Supplementary Figures 1-4, Supplementary Tables 1, 2, 4, 5, 7-11 for "Low Abundance of Circulating Tumor DNA in Localized Prostate Cancer"

#### SUPPLEMENTARY INFORMATION

- **SUPPLEMENTARY TABLES**
- **SUPPLEMENTARY FIGURES**

**SUPPLEMENTARY TABLES**

**Supplementary Table 1**

| *Gene;Allele* | *Direction* | *Sequence (5′ → 3′)* | *Amplicon Size (bp)* |
|---|---|---|---|
| CPSF3L;g.chr1:1254841C>G | Forward | CTTTGATCATCTGGGAGGTGAAGAA | 139 |
| | Reverse | CGAGATGGTGGGCTACGAC | |
| ZEB1;g.chr10:31810782A>C | Forward | CCATTGCTGACCAGAACAGTGT | 139 |
| | Reverse | CTGGATCACTTTCAAGGGTGGTT | |
| SORL1;g.chr11:121475922T>A | Forward | GATGAGTTGACTGTGTACAAAGTACAGA | 139 |
| | Reverse | GATGGGACCTACCTGTAGTAGACATTA | |
| CAMK2A;g.chr5:149644597T>G | Forward | CACAAGGCTCACCGATGTTG | 132 |
| | Reverse | GGGAGATGGAGGGTAATGGTTC | |
| TMEM140;g.chr7:134849280T>A | Forward | AGCTGCTGTTCATGAGCATCAT | 139 |
| | Reverse | CTCATTCCACAGGCAGAAGTTATAGAAG | |
| FER1L6;g.chr8:125076589T>G | Forward | TGACGTATTTGAATCCGAATCTGTCTG | 139 |
| | Reverse | CAGAGGACTCAGTGGCTGTTAG | |
| SLC39A4;g.chr8:145639320G>C | Forward | TGTGAGGGTAGGTGCTCAGT | 138 |
| | Reverse | CGGGCTCTACGCCTTCTTC | |
| NOC2L;g.chr1:887560A>C | Forward | CATACTGCCAGTTGTACACAGACT | 138 |
| | Reverse | CCCGTGTGCAAGATGAGGAAAT | |

**Supplementary Table 1.** Locus specific primer sequences for the eight targets selected for spike-in experiments.

**Supplementary Table 2**

| *Approximate spike-in quantity* | $10^1$ | $10^2$ | $10^3$ | $10^4$ | $10^5$ | $10^6$ |
|---|---|---|---|---|---|---|
| *Gene;Allele* | Average unique mutant reads | | | | | |
| CPSF3L;g.chr1:1254841C>G | 32 | 79 | 299 | 686 | 1826 | 2396 |
| ZEB1;g.chr10:31810782A>C | 0 | 27 | 156 | 409 | 1101 | 1016 |
| SORL1;g.chr11:121475922T>A | 2 | 18 | 161 | 470 | 815 | 1167 |
| CAMK2A;g.chr5:149644597T>G | 1 | 19 | 122 | 404 | 772 | 1039 |
| TMEM140;g.chr7:134849280T>A | 2 | 6 | 55 | 118 | 143 | 114 |
| FER1L6;g.chr8:125076589T>G | 0 | 12 | 110 | 293 | 883 | 678 |
| SLC39A4;g.chr8:145639320G>C | 4 | 24 | 146 | 392 | 832 | 1172 |
| NOC2L;g.chr1:887560A>C | 13 | 12 | 92 | 211 | 427 | 951 |
| *Inter-target average* | 7 | 25 | 142 | 373 | 850 | 1067 |
| *Inter-target standard deviation* | 12.0 | 23.2 | 72.4 | 172.3 | 493.4 | 639.7 |
| *Detection rate* | 75% | 100% | 100% | 100% | 100% | 100% |

**Supplementary Table 2.** Average duplicate-marked read counts for mutant alleles from a spike-in experiment. Purified fragments of product corresponding to the region flanking each allele were spiked in approximate quantities into aliquots of healthy donor plasma (3 mL). Data represents the average of two independent experiments performed in duplicate with two different pools of synthetic DNA for a total of eight libraries per target. Locus-specific primer sequences for each target are given in Supplementary Table 1.

**Supplementary Table 3**

See Microsoft Excel file.

**Supplementary Table 3.** Catalog of somatic mutations from exome sequencing of plasma from patients with metastatic prostate cancer.

**Supplementary Table 4**

| *Case* | *Gene;Allele* | *Direction* | *Sequence (5′ → 3′)* | *Amplicon Size (bp)* |
|---|---|---|---|---|
| M03 | TP53;g.chr17:7578406C>T | Forward | CTAAGAGCAATCAGTGAGGAATCAGA | 139 |
| | | Reverse | GCACATGACGGAGGTTGTGA | |
| | LIM2;g.chr19:51885774C>T | Forward | GAAGGTAGGCTGATGAGCGAA | 135 |
| | | Reverse | GGAGGTGTGGGAATGACCATG | |
| | ZFHX4;g.chr8:77616607A>T | Forward | CAGCAAAGGAGATACCCTGCAA | 139 |
| | | Reverse | ACTGTCCTCTAACTCGCTGATCT | |
| | ZNF276;g.chr16:89789061G>C | Forward | ATGGAGAGGCCATCCGCAGA | 137 |
| | | Reverse | GCTGTGGCACTGGTAGAACTG | |
| | MYH7; g.chr14:23884954_23884955ins | Forward | CCCGCTCACTAGTCTCAATCAG | 139 |
| | | Reverse | CCTGAAGGAGAACATCGCCAT | |
| | COG8;g.chr16:69368813G>T | Forward | GACAGCCCAAAGTACATGCAC | 139 |
| | | Reverse | GTGAATGAGAGTGCCATCTTCCAT | |
| | NSG2;g.chr5:173473835C>T | Forward | TAGGTCTGGGAAGAAAGGCGTA | 139 |
| | | Reverse | GCAGCTGTAAGTGATTAACCTCCAAG | |
| M04 | TM6SF2;g.chr19:19375665C>A | Forward | CATACAGCAGATTGCACACGAA | 137 |
| | | Reverse | CCTGACATCCATCTTCCCTGTCT | |
| | PABPC1;g.chr8:101721935G>C | Forward | CGTTCATTTCTGTAACTGCTTTAGTGG | 139 |
| | | Reverse | GCTCCTCATTCTACCATTACGTTAACTTTTT | |
| | PGLYRP3;g.chr1:153275064T>A | Forward | CTGGGAGGTTCATTTTAGGGCA | 139 |
| | | Reverse | GAGTTGAGTGATTGCTCACAGAGA | |
| | TRAF5;g.chr1:211527783G>C | Forward | CTGGTCTTGGAAATGCTGTTGAG | 139 |
| | | Reverse | AGCAGTGACTTTCTTACCTCCTGA | |
| | GRID1;g.chr10:87614319A>T | Forward | TGTCTTTGCCAAAAGTCTCACTATAGG | 139 |
| | | Reverse | TCCAATGGGTTTTTCTCCTCTTCTG | |
| | ATP2B2;g.chr3:10381924A>G | Forward | GGTCCTGGATTCTCCATCCAGT | 129 |
| | | Reverse | GACCAGTGGATGTGGTGCATAT | |
| | FRG1;g.chr4:190878562G>C | Forward | AGGGAGGAAAAATTAATTCAGTCTAAACACT | 139 |
| | | Reverse | CTTCATTGCATCTAATAAAGCAGCTATTTGA | |
| M06 | APC; g.chr5:112175951_112175952ins | Forward | GAATCAGAGCAGCCTAAAGAATCAAATG | 139 |
| | | Reverse | GGCATGGCAGAAATAATACATTCTTCTAGT | |
| | ORC5;g.chr7:103801558C>A | Forward | CTGAAGAATCCCACAAAGATGACAGA | 139 |
| | | Reverse | GCCAAAACCATTTCCACTAGACAGATTATT | |
| | SPO11;g.chr20:55915197A>C | Forward | GCTAGTTTTCTACAATTTAGCCTCACCTA | 135 |
| | | Reverse | TCTAACCTTTTAAGATCAGAAGGGAGAAGA | |
| | LRP1B;g.chr2:141607694C>A | Forward | GTTGTAACATGTACGAATCTTTTACCTCTT | 139 |
| | | Reverse | ACTGTGATAGACTTCGATGCATCTG | |
| | CASP1;g.chr11:104901134C>A | Forward | TGTATAATAGAACTTACCGAAGCAGTGAGA | 139 |
| | | Reverse | TGCAATGAAGAATTTGACAGTATTCCTAGA | |
| | NIPSNAP1;g.chr22:29965251G>C | Forward | AGTGAGCATGGGTAGTCCTCAT | 138 |
| | | Reverse | CAGGGTCTGTTTGAACTGAGTTTG | |
| | OVCH1;g.chr12:29642561C>A | Forward | CACCCACATGTTACTGAACAGCTA | 139 |
| | | Reverse | CGTGTAATACTGTGCTCAAGAGCAT | |
| | THBS1;g.chr15:39879635G>C | Forward | CGAGCTGTGGCAATGGAATTC | 139 |
| | | Reverse | CCCTGAGAGGCTAAGATGCTTAC | |

| | | | | |
|---|---|---|---|---|
| | MSL1;g.chr17:38285608C>A | Forward | GGAAGTACAGAAAGAAAGACTCCTGT | 139 |
| | | Reverse | CTCAATTCACGTTTACAAACAGTTTCAGAT | |
| | TDRD12;g.chr19:33288751T>G | Forward | GGGTGGAAGAAGGCCCAATTTA | 129 |
| | | Reverse | CCCTTGGAAGCTTTGTATTTTTGGC | |
| | RCHY1;g.chr4:76434490T>A | Forward | CTTCACTTTAAAGCGATCTAGTTGATGATC | 139 |
| | | Reverse | GCACACTTAGTTTCTTTTGCTTCTTACT | |
| | MCM4;g.chr8:48874964C>A | Forward | GAAGATATAGTGGCAAGTGAGCAGT | 139 |
| | | Reverse | CAACATTTTCAGAGCCCAAAGTAAGAATTT | |
| M07 | LAMA2;g.chr6:129813480C>A | Forward | GATTGGGCTTAATTGTAAGGAAGCAA | 139 |
| | | Reverse | CGGAGTTTCTGATGGGCACAG | |
| | DHX9;g.chr1:182811710C>G | Forward | TCCTCCCAGTTTTTGCTATTATAAATGCT | 132 |
| | | Reverse | GGGTCATCTTCCTTTTGCCACA | |
| | MAGEA4;g.chrX:151092687G>T | Forward | CCTCCGAGTCCCTGAAGATGAT | 135 |
| | | Reverse | GTCTTGGGAAAGATCTGATTATTACCCAG | |
| | CES1;g.chr16:55855305G>T | Forward | CCATAGCTGGATTGACAGGTGT | 136 |
| | | Reverse | GCTCTGTGACCATCTTTGGAGA | |
| | MFSD2A;g.chr1:40433489T>A | Forward | CTGGGTTGAGTTGGGATGTCT | 139 |
| | | Reverse | CACAGCTACCGCATATGTAATGATGA | |
| | S100A7A;g.chr1:153391713G>T | Forward | CTTCACAGGACAAAAAGGGCATAC | 139 |
| | | Reverse | ATGGCTCTGCTTGTGGTAGTC | |
| | HARBI1;g.chr11:46637383G>T | Forward | TCGGTTCACATAGGAGAGGTCTT | 139 |
| | | Reverse | CAGCTGATGAAGCCTCCATTCA | |
| | TMEM200A;g.chr6:130762831T>G | Forward | CACACTCAAAGTCCTTGGACTTAGA | 139 |
| | | Reverse | CGGGTCTTCTTTGTTCTCTAGTTTCATAT | |
| | PDE1C;g.chr7:31918688T>G | Forward | ACTGCGTGAACGATGCTCTT | 139 |
| | | Reverse | GATGAGCTCAGTGACATTCAGTCA | |
| | DIAPH2;g.chrX:96328009G>T | Forward | GCAGAAGCATTAGAAGAAAAGAAGACTG | 138 |
| | | Reverse | CCCTCTCACCGTTACCTATACCAAT | |

**Supplementary Table 4.** Locus specific primer sequences for the ctDNA targets in patients with metastatic prostate cancer.

**Supplementary Table 5**

| *Case* | *ERG status* | *Pathologic feature* | *Sequence pool* | *% of pool* |
|---|---|---|---|---|
| L001 | Positive | Gp3 | L001-1 | 18.6% |
| | Positive | Gp3 | L001-1 | 58.9% |
| | Positive | Gp4 | L001-1 | 22.5% |
| | Positive | Gp3 | L001-2 | 50.0% |
| | Positive | Gp4 | L001-2 | 50.0% |
| L002 | Negative | Gp3 | L002-1 | 33.3% |
| | Negative | Gp4 | L002-1 | 33.3% |
| | Negative | Gp4 | L002-1 | 33.3% |
| | Negative | Gp3 | L002-2 | 10.0% |
| | Negative | Gp3 | L002-2 | 10.0% |
| | Negative | Gp4 | L002-2 | 80.0% |
| L003 | Positive | Gp3 | L003-1 | 26.7% |
| | Positive | Gp4 | L003-1 | 73.3% |
| | Positive | Gp3 | L003-2 | 50.0% |
| | Positive | Gp4 | L003-2 | 50.0% |
| L004 | Negative | Gp3 | L004-1 | 100% |
| | Negative | Gp3 | L004-2 | 100% |
| | Negative | Gp4 | L004-3 | 100% |
| | Negative | HGPIN | L004-4 | 100% |
| | Negative | HGPIN | L004-5 | 100% |
| | Negative | HGPIN | L004-6 | 100% |
| | Negative | HGPIN | L004-7 | 100% |
| | Negative | HGPIN | L004-8 | 100% |
| | Negative | HGPIN | L004-9 | 100% |
| L015 | Positive | Gp5 | L015-1 | 100% |
| | Positive | IDC | L015-2 | 100% |
| | Positive | Gp5 | L015-3 | 100% |
| | Positive | IDC | L015-4 | 100% |
| | Positive | Gp5 | L015-5 | 100% |
| L039 | Negative | Gp5 | L039-1 | 100% |
| | Negative | Gp5 | L039-2 | 100% |
| | Negative | Gp5 | L039-3 | 100% |
| | Negative | IDC | L039-4 | 100% |
| | Negative | IDC | L039-5 | 100% |
| | Negative | Gp4 | L039-6 | 51.8% |
| | Negative | Gp4 | L039-6 | 48.2% |
| | Negative | Gp4 | L039-7 | 45.9% |
| | Negative | Gp4 | L039-7 | 54.1% |
| | Negative | Gp5 | L039-8 | 100% |
| | Negative | Gp5 | L039-9 | 100% |
| | Negative | IDC | L039-10 | 100% |

| | | | | |
|---|---|---|---|---|
| | Negative | Lymph Node | L039-11 | 100% |
| L040 | Negative | Gp3 | L040-1 | 100% |
| | Negative | Gp3 | L040-2 | 100% |
| | Negative | Gp3 | L040-3 | 100% |
| | Negative | Gp4 | L040-4 | 100% |
| | Negative | Gp4 | L040-5 | 100% |
| | Negative | Gp4 | L040-6 | 100% |
| | Negative | HGPIN | L040-7 | 100% |
| L107 | Positive | Gp3 | L107-1 | 13.8% |
| | Positive | Gp3 | L107-1 | 86.2% |
| | Positive | HGPIN | L107-2 | 100% |
| | Positive | Gp3 | L107-3 | 1.1% |
| | Positive | Gp3 | L107-3 | 98.9% |
| | Positive | Gp3 | L107-4 | 15.0% |
| | Positive | Gp3 | L107-4 | 85.0% |
| L108 | Negative | Gp3 | L108-1 | 47.5% |
| | Negative | Gp3 | L108-1 | 52.5% |
| | Negative | IDC | L108-2 | 47.1% |
| | Negative | IDC | L108-2 | 52.9% |
| | Negative | Gp3 | L108-3 | 49.3% |
| | Negative | Gp3 | L108-3 | 50.7% |
| | Negative | IDC | L108-4 | 100% |
| | Negative | Gp3 | L108-5 | 41.6% |
| | Negative | Gp3 | L108-5 | 58.4% |
| | Negative | Gp4 | L108-6 | 49.3% |
| | Negative | Gp4 | L108-6 | 50.7% |
| | Negative | HGPIN | L108-7 | 61.1% |
| | Negative | HGPIN | L108-7 | 38.9% |
| | Negative | Gp4 | L108-8 | 28.1% |
| | Negative | Gp4 | L108-8 | 71.9% |

**Supplementary Table 5.** Listing of each neoplastic histologic feature laser capture microdissected and sequenced for the identification of mutations to detect in circulating tumor DNA from matched plasma.

**Supplementary Table 6**

See Microsoft Excel file.

**Supplementary Table 6.** Catalog of somatic mutations from exome sequencing of tissue from patients with localized prostate cancer.

**Supplementary Table 7**

| *Case* | *Mutation* | *Classification* | *Focus/Foci harboring mutation* |
|---|---|---|---|
| L001 | g.chr1:236132293A>C | Trunk | L001-1, L001-2 |
| | g.chr11:126418270C>A | Trunk | L001-1, L001-2 |
| | g.chr11:58140876T>A | Trunk | L001-1, L001-2 |
| | g.chr12:123824014T>A | Branch | L001-1 |
| | g.chr14:96150262G>C | Trunk | L001-1, L001-2 |
| | g.chr4:155601476C>A | Branch | L001-1 |
| | g.chr4:187115632C>A | Trunk | L001-1, L001-2 |
| | g.chr7:143052765A>C | Branch | L001-2 |
| | g.chr7:1607899G>C | Trunk | L001-1, L001-2 |
| | g.chr8:25231044T>G | Trunk | L001-1, L001-2 |
| | g.chr9:117096905C>A | Trunk | L001-1, L001-2 |
| L002 | g.chr1:116836011G>T | Trunk | L002-1, L002-2 |
| | g.chr1:39816681G>C | Trunk | L002-1, L002-2 |
| | g.chr1:42182700T>G | Branch | L002-2 |
| | g.chr12:131745917C>A | Branch | L002-1 |
| | g.chr19:7104596C>G | Trunk | L002-1, L002-2 |
| | g.chr22:46199148A>C | Trunk | L002-1, L002-2 |
| | g.chr5:149026448T>G | Trunk | L002-1, L002-2 |
| | g.chr6:3526324G>T | Branch | L002-1 |
| | g.chr7:6779602T>G | Branch | L002-2 |
| | g.chr8:48517143C>G | Trunk | L002-1, L002-2 |
| L003 | g.chr15:75648676G>T | Trunk | L003-1, L003-2 |
| | g.chr17:78426705G>T | Branch | L003-1 |
| | g.chr19:28871188G>T | Branch | L003-2 |
| | g.chr19:9089900C>A | Trunk | L003-1, L003-2 |
| | g.chr5:169135903G>C | Trunk | L003-1, L003-2 |
| | g.chr6:3546468G>C | Trunk | L003-1, L003-2 |
| | g.chr7:30516535G>C | Trunk | L003-1, L003-2 |
| | g.chr9:135079069T>G | Trunk | L003-1, L003-2 |
| L004 | g.chr1:151162479_151162485del | Branch | L004-3, L004-8, L004-9 |
| | g.chr1:3789073C>G | Leaf | L004-2 |
| | g.chr1:6291991C>A | Leaf | L004-2 |
| | g.chr22:19220788G>C | Branch | L004-3, L004-4, L004-6 |
| | g.chr6:36465642C>A | Leaf | L004-2 |
| | g.chr8:110534476A>T | Leaf | L004-3 |
| L015 | g.chr14:77698059C>G | Branch | L015-1, L015-3, L015-5 |
| | g.chr17:4905798C>A | Branch | L015-1, L015-3, L015-4, L015-5 |
| | g.chr19:19337554G>T | Leaf | L015-2 |
| | g.chr19:49671565T>G | Trunk | L015-1, L015-2, L015-3, L015-4, L015-5 |
| | g.chr20:31024908A>T | Leaf | L015-1 |
| | g.chr20:37555195A>T | Branch | L015-1, L015-3, L015-4, L015-5 |
| | g.chr21:37759995C>A | Leaf | L015-1 |
| | g.chr3:160156803G>T | Leaf | L015-1 |
| | g.chr5:179201100G>T | Branch | L015-3, L015-5 |
| | g.chr6:26027236_26027237del | Branch | L015-3, L015-5 |
| | g.chr9:86588276C>A | Leaf | L015-3 |
| L039 | g.chr11:104821794G>A | Branch | L039-4, L039-5 |

| | | | |
|---|---|---|---|
| | g.chr12:28116381G>A | Branch | L039-1, L039-2 |
| | g.chr12:49218069G>A | Branch | L039-1, L039-2 |
| | g.chr14:75138135_75138136del | Leaf | L039-5 |
| | g.chr14:94844884C>T | Branch | L039-8, L039-10 |
| | g.chr3:50294284_50294367del | Branch | L039-5, L039-7 |
| | g.chr5:6609940T>C | Branch | L039-8, L039-10 |
| | g.chr6:53764594G>A | Branch | L039-4, L039-8, L039-10 |
| L040 | g.chr1:3428160T>G | Branch | L040-1, L040-2, L040-3, L040-4 |
| | g.chr10:50944443T>A | Branch | L040-6, L040-7 |
| | g.chr10:69959242C>A | Branch | L040-1, L040-2, L040-4, L040-7 |
| | g.chr19:17330030C>A | Leaf | L040-3 |
| | g.chr19:41383849C>G | Leaf | L040-4 |
| | g.chr2:230861519G>T | Branch | L040-4, L040-5 |
| | g.chr20:2290333C>A | Branch | L040-3, L040-4, L040-5 |
| | g.chr3:132166224C>A | Leaf | L040-1 |
| | g.chr6:32610403C>A | Branch | L040-2, L040-4, L040-5 |
| | g.chr7:55238874T>A | Branch | L040-4, L040-5 |
| | g.chr9:32459450T>G | Branch | L040-1, L040-4, L040-5 |
| L107 | g.chr1:152191717A>T | Leaf | L107-3 |
| | g.chr10:69934258C>G | Branch | L107-2, L107-3, L107-4 |
| | g.chr11:128333503T>C | Branch | L107-2, L107-3, L107-4 |
| | g.chr11:60102507G>T | Trunk | L107-1, L107-2, L107-3, L107-4 |
| | g.chr16:89212430C>T | Leaf | L107-3 |
| L108 | g.chr1:11561278G>C | Leaf | L108-4 |
| | g.chr1:57173271C>G | Leaf | L108-2 |
| | g.chr11:36597104C>G | Branch | L108-3, L108-4, L108-6 |
| | g.chr13:58208018C>T | Branch | L108-3, L108-4, L108-6, L108-8 |
| | g.chr16:30977694A>G | Branch | L108-3, L108-4, L108-6, L108-8 |
| | g.chr17:17881032C>A | Branch | L108-6, L108-8 |
| | g.chr17:6023871C>A | Leaf | L108-2 |
| | g.chr2:33412052A>T | Leaf | L108-1 |
| | g.chr20:19956215C>T | Branch | L108-1, L108-2 |
| | g.chr6:31797631C>T | Branch | L108-3, L108-4, L108-6, L108-8 |
| | g.chr7:117232188A>G | Leaf | L108-4 |
| | g.chr8:22548872G>T | Branch | L108-3, L108-4, L108-6, L108-8 |
| | g.chr9:135458585C>T | Leaf | L108-6 |

**Supplementary Table 7.** Point mutations discovered in tissue selected for analysis in plasma. The basis for the selection of each mutation is shown with respect to foci harboring each mutation. Locus-specific primer sequences for each target are given in Supplementary Table 8.

**Supplementary Table 8**

| *Case* | *Gene;Allele* | *Direction* | *Sequence (5′ → 3′)* | *Amplicon Size (bp)* |
|---|---|---|---|---|
| L001 | g.chr1:236132293A>C | Forward | CTCTCTCATGGTCTGAAATTGACTGG | 137 |
| | | Reverse | CTCAGCCTACAGCAGACATCAA | |
| | g.chr11:126418270C>A | Forward | CCTTCTCTTGAGGGCTTTCTCTTT | 134 |
| | | Reverse | GTATTTCTGCACCAGCTCCTGTA | |
| | g.chr11:58140876T>A | Forward | AGAAAATAGATCCATTCTTCCCTCTACTCA | 139 |
| | | Reverse | AATTTGAAGCAAAACCACCTTAGTACAC | |
| | g.chr12:123824014T>A | Forward | GCCTACTCCTACTCATCCCGAA | 139 |
| | | Reverse | TGAGTTTGAAGGACTTTGGACTTAGTCTA | |
| | g.chr14:96150262G>C | Forward | GCCATTTTGGCCCAGTACCTA | 128 |
| | | Reverse | CCCACTTGCCGGAAGGATTC | |
| | g.chr4:155601476C>A | Forward | AAATAGGGTGACTGTGATTTGAGTAACTTT | 138 |
| | | Reverse | CTTGTGGCCAGTCCCATATAACA | |
| | g.chr4:187115632C>A | Forward | ATCAAAATGAGAAAACAAACCTTTGTCCA | 129 |
| | | Reverse | GTGGCGGTATTCCTCTGTGTAC | |
| | g.chr7:143052765A>C | Forward | TGCATCCTGCGTTGGTGTAT | 137 |
| | | Reverse | CCTAAGGATCTGGGAAAAGACCCA | |
| | g.chr7:1607899G>C | Forward | CAAAACTGCTGCTTCTGCTTGT | 139 |
| | | Reverse | CATCTGGCGGACAAATACCC | |
| | g.chr8:25231044T>G | Forward | GGGATATTTGTTAAAGGGAAACAGTGATC | 126 |
| | | Reverse | AAAAAGGATCACAGAATCCCAACATAAATC | |
| | g.chr9:117096905C>A | Forward | TCCCTGGAAATCAGCTGACATC | 139 |
| | | Reverse | AACAAGAGGCTGTCAATCACGA | |
| L002 | g.chr1:116836011G>T | Forward | GGGATCTAGGAGGTATGAGAGATACTG | 138 |
| | | Reverse | GTCCTCTAGAGGTTCACAGCCTA | |
| | g.chr1:39816681G>C | Forward | TTCATCAGGCTAAGGAGCAATATGAG | 139 |
| | | Reverse | TGAAAAATATCAAGCGTCATAGCAGTCTAT | |
| | g.chr1:42182700T>G | Forward | GTTGTGGAAGACGAGTATGTTTACTCA | 139 |
| | | Reverse | TGGGAAACCAAGGTCACAACTG | |
| | g.chr12:131745917C>A | Forward | CAGTCCTGGGAGATGTGTGAAT | 139 |
| | | Reverse | AGAAACCTCAAGCTCCGATGATG | |
| | g.chr19:7104596C>G | Forward | TCCGCTACCTCCACCTCAATAT | 139 |
| | | Reverse | TGGATTCTGTAGTGGATGCTGTAGT | |
| | g.chr22:46199148A>C | Forward | CCCAAGGCCAACAGATCTCATA | 139 |
| | | Reverse | GTATGTGCAAAAGCTGGAAGATAAGATG | |
| | g.chr5:149026448T>G | Forward | AAGACAGGCTTTCAAAGCCAGA | 128 |
| | | Reverse | CCGCGTCTTAGAAGTCAGTGA | |
| | g.chr6:3526324G>T | Forward | CTCCCGTGTTTTACCACTTCAGT | 139 |
| | | Reverse | AACACGTCTGCACTAGCTTTCA | |
| | g.chr7:6779602T>G | Forward | GAAACAACAATATCAGCTCCTTTCTCTTTT | 139 |
| | | Reverse | GAAACGAAGGCAGAAGAATCGTT | |
| | g.chr8:48517143C>G | Forward | GAAGCAGCTACTGCCGTAGT | 129 |
| | | Reverse | CCCAACCAATCCATTATGAAGCCAT | |
| L003 | g.chr15:75648676G>T | Forward | CCTGCAGAGGAACACGTCT | 137 |
| | | Reverse | CCTGGAGTGCGTTTTCCGT | |
| | g.chr17:78426705G>T | Forward | TTCACGGCCACAAAGAGCTCAA | 139 |
| | | Reverse | GGCTCTAGATGTCAAGCTCACATTTAC | |
| | g.chr19:28871188G>T | Forward | GCCATCAACCCAGGATTCGAAG | 139 |
| | | Reverse | TGGACTCCGAATACAGCTGTTATG | |
| | g.chr19:9089900C>A | Forward | CGTTCGTGGCCAGAGTCAAA | 129 |
| | | Reverse | TTTGCTGTTCCCACTGGGATT | |
| | g.chr5:169135903G>C | Forward | CAGCCTTCTTCCTTTTAATGAGAGC | 139 |

| | | | | |
|---|---|---|---|---|
| | | Reverse | CACAGAGCAGGTGAATGACTTG | |
| | g.chr6:3546468G>C | Forward | CAGATTCCAGTCTGGGTCTGAAA | 139 |
| | | Reverse | GAGGCATTGAGAGCCTCAGTAG | |
| | g.chr7:30516535G>C | Forward | ACATTCTCCCTCATAGTGGCTGTA | 139 |
| | | Reverse | TTCAGCGAGATGATGTGCATCA | |
| | g.chr9:135079069T>G | Forward | TCAAAAAGTTGTTTAAACTCTGTGCAA | 139 |
| | | Reverse | CCCATCGAATGCAGCTGCTA | |
| L004 | g.chr1:151162479_151162485del | Forward | TCCCTTCAGGGCTCCTCTTT | 130 |
| | | Reverse | ACCGGCTTTCTGGGCTTTT | |
| | g.chr1:3789073C>G | Forward | GCCTTCCCTCATTGTCTTTTGG | 139 |
| | | Reverse | GTGAGCTTACATGTGATCCAGGTT | |
| | g.chr1:6291991C>A | Forward | CCTGAGTTCCAATACTCCAGTAAAACC | 139 |
| | | Reverse | GTTTACAGCTGGCTCCAATTTCAAC | |
| | g.chr22:19220788G>C | Forward | ACGTTACCTTTTTGGCATAGAGCA | 139 |
| | | Reverse | CGGAGACTTGGTCAAAACCACT | |
| | g.chr6:36465642C>A | Forward | TCCCAGTCAACGCCTTCAAAA | 139 |
| | | Reverse | CTTTTTCTGAGGAGACTGCTTGAGAT | |
| | g.chr8:110534476A>T | Forward | TGAACTCCAGGAAATTGCTGGT | 139 |
| | | Reverse | TTACCTTAATCCACCCAGAGTCACT | |
| L015 | g.chr14:77698059C>G | Forward | GGACGACGCACGCATCTA | 128 |
| | | Reverse | GAGCTTTGCCTACCCAGAACA | |
| | g.chr17:4905798C>A | Forward | CAGGTGAGCTATATGGAGATCTACTGT | 139 |
| | | Reverse | TAGGAGGTCACAGCCAATTTGG | |
| | g.chr19:19337554G>T | Forward | AGGAGTCTCAACAGACCCTCA | 139 |
| | | Reverse | CTCCGTGTGTGAACTTGCTG | |
| | g.chr19:49671565T>G | Forward | GTCTCTGTCTTATTCTTTGTTTCTCTCCC | 139 |
| | | Reverse | CCATCTGATGGTCCCGTACA | |
| | g.chr20:31024908A>T | Forward | CCAACTGAGTTCCACCAGCTTT | 139 |
| | | Reverse | TGAACATTTGCAAGGAAAGTGATGC | |
| | g.chr20:37555195A>T | Forward | CCGAGCTGTTCAGCGAGTC | 129 |
| | | Reverse | ATGGCGTGCACCTCATCGG | |
| | g.chr21:37759995C>A | Forward | GGTAGAAAAGGGACCAGATGGAAA | 139 |
| | | Reverse | GACAGCAATACTCACCATCTCCT | |
| | g.chr3:160156803G>T | Forward | AATAGCCCTTAGTGCAAAATTAACAGGTA | 139 |
| | | Reverse | ACATTCTTCAGGCATCTGGTAACTTTTAT | |
| | g.chr5:179201100G>T | Forward | CCTGGACATGCTTCAGTTTCCTC | 139 |
| | | Reverse | TGGGCAACTATCTGGGTTATGC | |
| | g.chr6:26027236_26027237del | Forward | CGTACAGAGTGCGTCCTTGA | 138 |
| | | Reverse | GAGACTCGTGGCGTTCTCAA | |
| | g.chr9:86588276C>A | Forward | ACCTCTCGAAGTTCTTTGATTTTAGCA | 138 |
| | | Reverse | ATTTTATAATTCTCCCTCCACAGTACCAAC | |
| L039 | g.chr11:104821794G>A | Forward | TCAAGTAGCTCCTTCATCCCTGT | 139 |
| | | Reverse | CATATCCTGCAGATCTATCCAATAAAGGAG | |
| | g.chr12:28116381G>A | Forward | CTCCCAGTCACTCCAGAGTCTAA | 139 |
| | | Reverse | AACAAGGTGGAGACGTACAAAGAG | |
| | g.chr12:49218069G>A | Forward | CCCAGAAGTACAGCAATGACTG | 130 |
| | | Reverse | CAACTCTCACCTGGCCTTCTG | |
| | g.chr14:75138135_75138136del | Forward | AAAATTTCAAAACTGACCTTCTTCATCCT | 137 |
| | | Reverse | ACCAGCCAGTGTTCTGACAATT | |
| | g.chr14:94844884C>T | Forward | GCGAGAGGCAGTTATTTTTGGG | 138 |
| | | Reverse | TAGAGGCCATACCCATGTCTATCC | |
| | g.chr3:50294284_50294367del | Forward | CATCATCTTCTGCGTAGCCTTGA | 139 |
| | | Reverse | GGTAGAATAGCAAGGTCAACGCTA | |
| | g.chr5:6609940T>C | Forward | CAAGAAGCTCCTCTTTCCTCTGT | 139 |
| | | Reverse | AAATCTTCAATGCCGTGGAATAAACG | |

| | | | | |
|---|---|---|---|---|
| | g.chr6:53764594G>A | Forward | GCACTGTGGCAGGTTTAAGTGT | 139 |
| | | Reverse | GAAGTCAGGCCACTGATTTCTTCA | |
| L040 | g.chr1:3428160T>G | Forward | GATGCAGGTCCTCTGATCTGTG | 139 |
| | | Reverse | CAGGGATTGAGATGGAAATCGTGAA | |
| | g.chr10:50944443T>A | Forward | GCTTGCCCATCACTTGACTGTA | 127 |
| | | Reverse | CTGCACGGGAAAGGTGTACTAT | |
| | g.chr10:69959242C>A | Forward | GGAGTCCACTCTCTGCTCATTG | 134 |
| | | Reverse | GTCCCAGCAGCAAACCTTACCT | |
| | g.chr19:17330030C>A | Forward | CCCACTCCATCTCTCATGTCCT | 138 |
| | | Reverse | CAGCTTTTCCTGGAGGTTCTGAT | |
| | g.chr19:41383849C>G | Forward | GGTGCTTCATGAGCAGCAAGA | 139 |
| | | Reverse | CTCCCTGCAGGAGGAGAAGAAC | |
| | g.chr2:230861519G>T | Forward | GATAACCATGTTGGCTTCTTTGAATTACAG | 139 |
| | | Reverse | GATAGTCAGGAGCAAATCGTCTGAG | |
| | g.chr20:2290333C>A | Forward | CACAGCTCTAGGAGTCCAGAGT | 139 |
| | | Reverse | GGCCTTTGTTCATGATCATTAAGACCT | |
| | g.chr3:132166224C>A | Forward | TAAGTGGCCTTATGGAGACATTTGC | 139 |
| | | Reverse | GAAGTTCTGTTCTGTGCTCTGTAGAAA | |
| | g.chr6:32610403C>A | Forward | TCACTTCCACAGAGCCTGAGAT | 128 |
| | | Reverse | GCAGGCCTTGGATGATGAAGAC | |
| L107 | g.chr1:152191717A>T | Forward | ACTCGTGTTGCCCAAAACCA | 139 |
| | | Reverse | TCCTAGCCGTGTCCGACAT | |
| | g.chr10:69934258C>G | Forward | CTTTCCATCCAAAATGAGCCACT | 139 |
| | | Reverse | TGAATCCGGCTGGTAGGAGA | |
| | g.chr11:128333503T>C | Forward | CCTCATCTGGGTCAGAAAGTTTGA | 139 |
| | | Reverse | ACAATGGCTTGTGTTTCTAGGCA | |
| | g.chr11:60102507G>T | Forward | GGATAGCCTGAAGAAACGTCTACAG | 135 |
| | | Reverse | CACACTCACAAGGATGACCAGT | |
| | g.chr16:89212430C>T | Forward | CTCAAACTGTTCTTCTATCCGCAGAT | 139 |
| | | Reverse | CTACCTGGCCCACTCTTTGAG | |
| L108 | g.chr1:11561278G>C | Forward | GCTTCGGGCTTCTGGAGTAC | 131 |
| | | Reverse | GTCCAGCGGTGGGTAGTAAAG | |
| | g.chr1:57173271C>G | Forward | AGATCTGGTTCCTCAACACCTCA | 139 |
| | | Reverse | GACAATGTGCTTCCGGTCAAAG | |
| | g.chr11:36597104C>G | Forward | TCAGTCTACATTTGTACTCTTTGTGATGC | 139 |
| | | Reverse | ACAGACTCATGGTAAGGGTTGGA | |
| | g.chr13:58208018C>T | Forward | ACTTCTACACGGTGGTGACTGA | 136 |
| | | Reverse | ACTTCTACACGGTGGTGACTGA | |
| | g.chr16:30977694A>G | Forward | CGAGACCTCAACCGCAAGATG | 138 |
| | | Reverse | GGCCCATTTCCTCCTGCAAA | |
| | g.chr17:17881032C>A | Forward | GTGATGGACGATGACATGCTCA | 139 |
| | | Reverse | CTTGGATACATACTCCGAAAGTCCA | |
| | g.chr17:6023871C>A | Forward | CCTTCCTCAACGTGTCTGTGA | 138 |
| | | Reverse | TAGCCATTGATCAAGTCTTTCATCTCC | |
| | g.chr2:33412052A>T | Forward | GTCAGTGCCCTCCAAATTTCAC | 139 |
| | | Reverse | AGGCAAGGTATGTGTTGAATGGAT | |
| | g.chr20:19956215C>T | Forward | GGGTGTTCAGCTCCTTCATG | 139 |
| | | Reverse | CACGTGGCACTCCTTGTTCT | |
| | g.chr6:31797631C>T | Forward | CATCAGCGGACTGTACCAGG | 138 |
| | | Reverse | AAAGACAAACATACTAAAGAACAAAGGCC | |
| | g.chr7:117232188A>G | Forward | GGTAGCAGCTATTTTTATGGGACATTTTC | 139 |
| | | Reverse | CGGTGTAAGGTCTCAGTTAGGATTG | |
| | g.chr8:22548872G>T | Forward | GCGCTCATGAGGCTAATGATGT | 133 |
| | | Reverse | CCCAACCCGGAACTCTCTTACT | |
| | g.chr9:135458585C>T | Forward | ACATCCTTGCCGACTGCAAA | 135 |

| | |
|---|---|
| Reverse | CCACTTTTGTCCAGCTTGTCTCT |

**Supplementary Table 8.** Locus specific primer sequences for the ctDNA targets in patients with localized prostate cancer.

**Supplementary Table 9**

| *Case* | *Mutation* | *Raw locus depth* | *Unique ref alleles* | *Unique alt alleles* | *Library read depth* |
|---|---|---|---|---|---|
| L001 | | | | | 954091 |
| | g.chr1:236132293A>C | 48301 | 16 | 0 | |
| | g.chr11:126418270C>A | 119355 | 30 | 0 | |
| | g.chr11:58140876T>A | 4916 | 4 | 0 | |
| | g.chr12:123824014T>A | 5065 | 5 | 0 | |
| | g.chr14:96150262G>C | 121864 | 30 | 0 | |
| | g.chr4:155601476C>A | 27910 | 15 | 0 | |
| | g.chr4:187115632C>A | 78955 | 15 | 0 | |
| | g.chr7:143052765A>C | 10394 | 5 | 0 | |
| | g.chr7:1607899G>C | 101173 | 22 | 0 | |
| | g.chr8:25231044T>G | 1881 | 5 | 0 | |
| | g.chr9:117096905C>A | 24466 | 9 | 0 | |
| L002 | | | | | 259916 |
| | g.chr1:116836011G>T | 21351 | 21 | 0 | |
| | g.chr1:39816681G>C | 9761 | 18 | 0 | |
| | g.chr1:42182700T>G | 9133 | 23 | 0 | |
| | g.chr12:131745917C>A | 781 | 8 | 0 | |
| | g.chr19:7104596C>G | 1161 | 23 | 0 | |
| | g.chr22:46199148A>C | 1889 | 10 | 0 | |
| | g.chr5:149026448T>G | 20441 | 31 | 0 | |
| | g.chr6:3526324G>T | 16902 | 22 | 0 | |
| | g.chr7:6779602T>G | 12983 | 17 | 0 | |
| | g.chr8:48517143C>G | 26479 | 28 | 0 | |
| L003 | | | | | 421763 |
| | g.chr15:75648676G>T | 36473 | 12 | 0 | |
| | g.chr17:78426705G>T | 12335 | 7 | 0 | |
| | g.chr19:28871188G>T | 13368 | 11 | 0 | |
| | g.chr19:9089900C>A | 51589 | 18 | 0 | |
| | g.chr5:169135903G>C | 35938 | 12 | 0 | |
| | g.chr6:3546468G>C | 24509 | 10 | 0 | |
| | g.chr7:30516535G>C | 29144 | 12 | 0 | |
| | g.chr9:135079069T>G | 24832 | 6 | 0 | |
| L004 | | | | | 790288 |
| | g.chr1:151162479_151162485del | 86872 | 20 | 0 | |
| | g.chr1:3789073C>G | 28470 | 10 | 0 | |
| | g.chr1:6291991C>A | 95334 | 22 | 0 | |
| | g.chr22:19220788G>C | 123292 | 29 | 0 | |
| | g.chr6:36465642C>A | 23033 | 11 | 0 | |
| | g.chr8:110534476A>T | 45857 | 9 | 0 | |
| L015 | | | | | 732729 |
| | g.chr14:77698059C>G | 45673 | 7 | 0 | |
| | g.chr17:4905798C>A | 32447 | 8 | 0 | |
| | g.chr19:19337554G>T | 21912 | 6 | 0 | |
| | g.chr19:49671565T>G | 70681 | 9 | 0 | |
| | g.chr20:31024908A>T | 13911 | 5 | 0 | |

| | | | | |
|---|---|---|---|---|
| | g.chr20:37555195A>T | 70506 | 12 | 0 | |
| | g.chr21:37759995C>A | 22981 | 7 | 0 | |
| | g.chr3:160156803G>T | 2517 | 2 | 0 | |
| | g.chr5:179201100G>T | 45673 | 13 | 0 | |
| | g.chr6:26027236_26027237del | 72856 | 11 | 0 | |
| | g.chr9:86588276C>A | 16891 | 5 | 0 | |
| L039 | | | | | 828342 |
| | g.chr11:104821794G>A | 23123 | 47 | 0 | |
| | g.chr12:28116381G>A | 31732 | 55 | 0 | |
| | g.chr12:49218069G>A | 83594 | 86 | 0 | |
| | g.chr14:75138135_75138136del | 50527 | 49 | 0 | |
| | g.chr14:94844884C>T | 65851 | 49 | 0 | |
| | g.chr3:50294284_50294367del | 152643 | 84 | 0 | |
| | g.chr5:6609940T>C | 52260 | 51 | 0 | |
| | g.chr6:53764594G>A | 52931 | 60 | 0 | |
| L040 | | | | | 733865 |
| | g.chr1:3428160T>G | 856 | 2 | 0 | |
| | g.chr10:50944443T>A | 19601 | 6 | 0 | |
| | g.chr10:69959242C>A | 180460 | 23 | 0 | |
| | g.chr19:17330030C>A | 26819 | 10 | 0 | |
| | g.chr19:41383849C>G | 3044 | 2 | 0 | |
| | g.chr2:230861519G>T | 5692 | 2 | 0 | |
| | g.chr20:2290333C>A | 83336 | 26 | 0 | |
| | g.chr3:132166224C>A | 1086 | 2 | 0 | |
| | g.chr6:32610403C>A | 25188 | 8 | 0 | |
| | g.chr7:55238874T>A | 50748 | 7 | 0 | |
| | g.chr9:32459450T>G | 5705 | 3 | 0 | |
| L107 | | | | | 1119431 |
| | g.chr1:152191717A>T | 107845 | 30 | 0 | |
| | g.chr10:69934258C>G | 41979 | 33 | 0 | |
| | g.chr11:128333503T>C | 43686 | 35 | 0 | |
| | g.chr11:60102507G>T | 108691 | 43 | 0 | |
| | g.chr16:89212430C>T | 198663 | 56 | 0 | |
| L108 | | | | | 418931 |
| | g.chr1:11561278G>C | 21151 | 106 | 0 | |
| | g.chr1:57173271C>G | 30260 | 99 | 0 | |
| | g.chr11:36597104C>G | 14554 | 70 | 0 | |
| | g.chr13:58208018C>T | 26259 | 99 | 0 | |
| | g.chr16:30977694A>G | 36949 | 94 | 0 | |
| | g.chr17:17881032C>A | 10319 | 103 | 0 | |
| | g.chr17:6023871C>A | 248 | 39 | 0 | |
| | g.chr2:33412052A>T | 17751 | 82 | 0 | |
| | g.chr20:19956215C>T | 39965 | 117 | 0 | |
| | g.chr6:31797631C>T | 21388 | 93 | 0 | |
| | g.chr7:117232188A>G | 4688 | 53 | 0 | |
| | g.chr8:22548872G>T | 25044 | 112 | 0 | |
| | g.chr9:135458585C>T | 95 | 15 | 0 | |

**Supplementary Table 9.** Read count data for bespoke cfDNA sequencing libraries from nine patients with localized prostate cancer, sampled from preoperative plasma. Reference and alternate reads count only unique molecules. Read depth: total number of mapped and unmapped reads.

**Supplementary Table 10**

| *Case* | *Days post-RP/Mutation* | *Raw locus depth* | *Unique ref alleles* | *Unique alt alleles* | *Library read depth* |
|---|---|---|---|---|---|
| L001 | 122 | | | | 725843 |
| | g.chr1:236132293A>C | 38566 | 23 | 0 | |
| | g.chr11:126418270C>A | 63800 | 32 | 0 | |
| | g.chr11:58140876T>A | 2102 | 5 | 0 | |
| | g.chr12:123824014T>A | 2973 | 5 | 0 | |
| | g.chr14:96150262G>C | 103754 | 53 | 0 | |
| | g.chr4:155601476C>A | 11842 | 15 | 0 | |
| | g.chr4:187115632C>A | 89253 | 31 | 0 | |
| | g.chr7:1430527655874A>C | 5874 | 9 | 0 | |
| | g.chr7:1607899G>C | 34886 | 18 | 0 | |
| | g.chr8:25231044T>G | 1188 | 6 | 0 | |
| | g.chr9:117096905C>A | 31336 | 19 | 0 | |
| | 234 | | | | 851599 |
| | g.chr1:236132293A>C | 85917 | 27 | 0 | |
| | g.chr11:126418270C>A | 50831 | 20 | 0 | |
| | g.chr11:58140876T>A | 5603 | 5 | 0 | |
| | g.chr12:123824014T>A | 3386 | 4 | 0 | |
| | g.chr14:96150262G>C | 123857 | 42 | 0 | |
| | g.chr4:155601476C>A | 11986 | 9 | 0 | |
| | g.chr4:187115632C>A | 103699 | 19 | 0 | |
| | g.chr7:143052765A>C | 4046 | 4 | 0 | |
| | g.chr7:1607899G>C | 56403 | 17 | 0 | |
| | g.chr8:25231044T>G | 1047 | 3 | 0 | |
| | g.chr9:117096905C>A | 25040 | 12 | 0 | |
| | 353 | | | | 884849 |
| | g.chr1:236132293A>C | 45012 | 14 | 0 | |
| | g.chr11:126418270C>A | 42173 | 14 | 0 | |
| | g.chr11:58140876T>A | 2457 | 3 | 0 | |
| | g.chr12:123824014T>A | 6899 | 6 | 0 | |
| | g.chr14:96150262G>C | 133419 | 52 | 0 | |
| | g.chr4:155601476C>A | 7432 | 7 | 0 | |
| | g.chr4:187115632C>A | 101795 | 26 | 0 | |
| | g.chr7:143052765A>C | 7058 | 7 | 0 | |
| | g.chr7:1607899G>C | 63064 | 24 | 0 | |
| | g.chr8:25231044T>G | 1036 | 4 | 0 | |
| | g.chr9:117096905C>A | 34031 | 16 | 0 | |
| | 472 | | | | 964686 |
| | g.chr1:236132293A>C | 77330 | 36 | 0 | |
| | g.chr11:126418270C>A | 68431 | 35 | 0 | |
| | g.chr11:58140876T>A | 1824 | 4 | 0 | |
| | g.chr12:123824014T>A | 2321 | 5 | 0 | |
| | g.chr14:96150262G>C | 136649 | 65 | 0 | |
| | g.chr4:155601476C>A | 16873 | 18 | 0 | |
| | g.chr4:187115632C>A | 57662 | 24 | 0 | |
| | g.chr7:143052765A>C | 3059 | 7 | 0 | |
| | g.chr7:1607899G>C | 97720 | 40 | 0 | |

| | | | | | |
|---|---|---|---|---|---|
| | g.chr8:25231044T>G | 598 | 4 | 0 | |
| | g.chr9:117096905C>A | 41935 | 26 | 0 | |
| L002 | 106 | | | | 1007266 |
| | g.chr1:116836011G>T | 71757 | 39 | 0 | |
| | g.chr1:39816681G>C | 43109 | 32 | 0 | |
| | g.chr1:42182700T>G | 44071 | 45 | 0 | |
| | g.chr12:131745917C>A | 2500 | 14 | 0 | |
| | g.chr19:7104596C>G | 6656 | 16 | 0 | |
| | g.chr22:46199148A>C | 6560 | 19 | 0 | |
| | g.chr5:149026448T>G | 41485 | 38 | 0 | |
| | g.chr6:3526324G>T | 60468 | 52 | 0 | |
| | g.chr7:6779602T>G | 56107 | 34 | 0 | |
| | g.chr8:48517143C>G | 105756 | 64 | 0 | |
| | 246 | | | | 678782 |
| | g.chr1:116836011G>T | 63843 | 19 | 0 | |
| | g.chr1:39816681G>C | 21167 | 12 | 0 | |
| | g.chr1:42182700T>G | 22354 | 16 | 0 | |
| | g.chr12:131745917C>A | 2341 | 5 | 0 | |
| | g.chr19:7104596C>G | 3336 | 14 | 0 | |
| | g.chr22:46199148A>C | 3760 | 6 | 0 | |
| | g.chr5:149026448T>G | 23224 | 20 | 0 | |
| | g.chr6:3526324G>T | 46748 | 18 | 0 | |
| | g.chr7:6779602T>G | 53207 | 18 | 0 | |
| | g.chr8:48517143C>G | 75706 | 28 | 0 | |
| L003 | 12 | | | | 579156 |
| | g.chr15:75648676G>T | 45497 | 24 | 0 | |
| | g.chr17:78426705G>T | 25280 | 16 | 0 | |
| | g.chr19:28871188G>T | 23755 | 22 | 0 | |
| | g.chr19:9089900C>A | 75874 | 31 | 0 | |
| | g.chr5:169135903G>C | 44679 | 16 | 0 | |
| | g.chr6:3546468G>C | 39505 | 18 | 0 | |
| | g.chr7:30516535G>C | 42577 | 18 | 0 | |
| | g.chr9:135079069T>G | 26780 | 11 | 0 | |
| | 96 | | | | 821136 |
| | g.chr15:75648676G>T | 39694 | 16 | 0 | |
| | g.chr17:78426705G>T | 62139 | 21 | 0 | |
| | g.chr19:28871188G>T | 46665 | 23 | 0 | |
| | g.chr19:9089900C>A | 78397 | 21 | 0 | |
| | g.chr5:169135903G>C | 52999 | 13 | 0 | |
| | g.chr6:3546468G>C | 58842 | 13 | 0 | |
| | g.chr7:30516535G>C | 51580 | 16 | 0 | |
| | g.chr9:135079069T>G | 72994 | 17 | 0 | |
| | 418 | | | | 833626 |
| | g.chr15:75648676G>T | 53516 | 21 | 0 | |
| | g.chr17:78426705G>T | 43535 | 19 | 0 | |
| | g.chr19:28871188G>T | 36946 | 25 | 0 | |
| | g.chr19:9089900C>A | 41034 | 15 | 0 | |
| | g.chr5:169135903G>C | 74530 | 22 | 0 | |
| | g.chr6:3546468G>C | 21587 | 12 | 0 | |
| | g.chr7:30516535G>C | 83208 | 28 | 0 | |

| | | | | | |
|---|---|---|---|---|---|
| | g.chr9:135079069T>G | 69205 | 18 | 0 | |
| | 572 | | | | 792204 |
| | g.chr15:75648676G>T | 43146 | 19 | 0 | |
| | g.chr17:78426705G>T | 39460 | 16 | 0 | |
| | g.chr19:28871188G>T | 33142 | 27 | 0 | |
| | g.chr19:9089900C>A | 73426 | 29 | 0 | |
| | g.chr5:169135903G>C | 65310 | 24 | 0 | |
| | g.chr6:3546468G>C | 59101 | 20 | 0 | |
| | g.chr7:30516535G>C | 50422 | 21 | 0 | |
| | g.chr9:135079069T>G | 88745 | 24 | 0 | |
| | 719 | | | | 995160 |
| | g.chr15:75648676G>T | 126828 | 22 | 0 | |
| | g.chr17:78426705G>T | 50412 | 11 | 0 | |
| | g.chr19:28871188G>T | 48793 | 13 | 0 | |
| | g.chr19:9089900C>A | 38202 | 8 | 0 | |
| | g.chr5:169135903G>C | 63278 | 11 | 0 | |
| | g.chr6:3546468G>C | 14998 | 6 | 0 | |
| | g.chr7:30516535G>C | 115828 | 19 | 0 | |
| | g.chr9:135079069T>G | 124793 | 13 | 0 | |
| L004 | 85 | | | | 734314 |
| | g.chr1:151162479_151162485del | 72046 | 17 | 0 | |
| | g.chr1:3789073C>G | 22375 | 8 | 0 | |
| | g.chr1:6291991C>A | 194497 | 40 | 0 | |
| | g.chr22:19220788G>C | 95687 | 24 | 0 | |
| | g.chr6:36465642C>A | 24981 | 9 | 0 | |
| | g.chr8:110534476A>T | 26188 | 10 | 0 | |
| | 120 | | | | 1007444 |
| | g.chr1:151162479_151162485del | 139447 | 25 | 0 | |
| | g.chr1:3789073C>G | 54668 | 16 | 0 | |
| | g.chr1:6291991C>A | 178848 | 29 | 0 | |
| | g.chr22:19220788G>C | 107370 | 22 | 0 | |
| | g.chr6:36465642C>A | 29385 | 10 | 0 | |
| | g.chr8:110534476A>T | 72070 | 17 | 0 | |
| | 295 | | | | 895012 |
| | g.chr1:151162479_151162485del | 170632 | 40 | 0 | |
| | g.chr1:3789073C>G | 27803 | 14 | 0 | |
| | g.chr1:6291991C>A | 129291 | 36 | 0 | |
| | g.chr22:19220788G>C | 116023 | 35 | 0 | |
| | g.chr6:36465642C>A | 19520 | 9 | 0 | |
| | g.chr8:110534476A>T | 52584 | 17 | 0 | |
| | 598 | | | | 976471 |
| | g.chr1:151162479_151162485del | 146245 | 35 | 0 | |
| | g.chr1:3789073C>G | 63594 | 22 | 0 | |
| | g.chr1:6291991C>A | 112071 | 29 | 0 | |
| | g.chr22:19220788G>C | 145724 | 39 | 0 | |
| | g.chr6:36465642C>A | 7079 | 4 | 0 | |
| | g.chr8:110534476A>T | 102733 | 31 | 0 | |
| L015 | 92 | | | | 1531476 |
| | g.chr14:77698059C>G | 31666 | 14 | 0 | |
| | g.chr17:4905798C>A | 72371 | 19 | 0 | |

| | | | | | |
|---|---|---|---|---|---|
| | g.chr19:19337554G>T | 47986 | 21 | 0 | |
| | g.chr19:49671565T>G | 160499 | 21 | 0 | |
| | g.chr20:31024908A>T | 16340 | 8 | 0 | |
| | g.chr20:37555195A>T | 214119 | 50 | 0 | |
| | g.chr21:37759995C>A | 6770 | 4 | 0 | |
| | g.chr3:160156803G>T | 6254 | 5 | 0 | |
| | g.chr5:179201100G>T | 181758 | 45 | 0 | |
| | g.chr6:26027237C>G | 65590 | 15 | 0 | |
| | g.chr9:86588276C>A | 17127 | 9 | 0 | |
| | 183 | | | | 593773 |
| | g.chr14:77698059C>G | 16503 | 7 | 0 | |
| | g.chr17:4905798C>A | 12211 | 5 | 0 | |
| | g.chr19:19337554G>T | 3339 | 3 | 0 | |
| | g.chr19:49671565T>G | 18330 | 4 | 0 | |
| | g.chr20:31024908A>T | 34665 | 8 | 0 | |
| | g.chr20:37555195A>T | 90831 | 13 | 0 | |
| | g.chr21:37759995C>A | 77432 | 9 | 0 | |
| | g.chr3:160156803G>T | 11340 | 4 | 0 | |
| | g.chr5:179201100G>T | 9861 | 4 | 0 | |
| | g.chr6:26027237C>G | 55785 | 9 | 0 | |
| | g.chr9:86588276C>A | 2561 | 1 | 0 | |
| | 239 | | | | 785887 |
| | g.chr14:77698059C>G | 20636 | 7 | 0 | |
| | g.chr17:4905798C>A | 18523 | 6 | 0 | |
| | g.chr19:19337554G>T | 15159 | 4 | 0 | |
| | g.chr19:49671565T>G | 43252 | 6 | 0 | |
| | g.chr20:31024908A>T | 80080 | 11 | 0 | |
| | g.chr20:37555195A>T | 63839 | 9 | 0 | |
| | g.chr21:37759995C>A | 0 | 0 | 0 | |
| | g.chr3:160156803G>T | 3211 | 2 | 0 | |
| | g.chr5:179201100G>T | 61140 | 9 | 0 | |
| | g.chr6:26027237C>G | 100564 | 4 | 0 | |
| | g.chr9:86588276C>A | 16800 | 3 | 0 | |
| L039 | 106 | | | | 573305 |
| | g.chr11:104821794G>A | 17708 | 40 | 0 | |
| | g.chr12:28116381G>A | 18645 | 48 | 0 | |
| | g.chr12:49218069G>A | 54232 | 76 | 0 | |
| | g.chr14:75138135_75138136del | 34322 | 51 | 0 | |
| | g.chr14:94844884C>T | 39077 | 43 | 0 | |
| | g.chr3:50294284_50294367del | 107727 | 66 | 0 | |
| | g.chr5:6609940T>C | 21757 | 39 | 0 | |
| | g.chr6:53764594G>A | 47599 | 50 | 0 | |
| | 169 | | | | 551245 |
| | g.chr11:104821794G>A | 22036 | 43 | 0 | |
| | g.chr12:28116381G>A | 16308 | 43 | 0 | |
| | g.chr12:49218069G>A | 64126 | 69 | 0 | |
| | g.chr14:75138135_75138136del | 27407 | 38 | 0 | |
| | g.chr14:94844884C>T | 41816 | 43 | 0 | |
| | g.chr3:50294284_50294367del | 115040 | 59 | 0 | |
| | g.chr5:6609940T>C | 27466 | 37 | 0 | |

| | | | | | |
|---|---|---|---|---|---|
| | g.chr6:53764594G>A | 55491 | 47 | 0 | |
| | 435 | | | | 528331 |
| | g.chr11:104821794G>A | 15329 | 41 | 0 | |
| | g.chr12:28116381G>A | 15814 | 47 | 0 | |
| | g.chr12:49218069G>A | 65666 | 83 | 0 | |
| | g.chr14:75138135_75138136del | 36808 | 48 | 0 | |
| | g.chr14:94844884C>T | 40765 | 44 | 0 | |
| | g.chr3:50294284_50294367del | 107287 | 84 | 0 | |
| | g.chr5:6609940T>C | 17499 | 37 | 0 | |
| | g.chr6:53764594G>A | 52032 | 53 | 0 | |
| L040 | 187 | | | | 933997 |
| | g.chr1:3428160T>G | 29489 | 14 | 0 | |
| | g.chr10:50944443T>A | 59096 | 18 | 0 | |
| | g.chr10:69959242C>A | 152578 | 19 | 0 | |
| | g.chr19:17330030C>A | 84854 | 31 | 0 | |
| | g.chr19:41383849C>G | 1739 | 3 | 0 | |
| | g.chr2:230861519G>T | 17928 | 6 | 0 | |
| | g.chr20:2290333C>A | 68232 | 18 | 0 | |
| | g.chr3:132166224C>A | 3743 | 4 | 0 | |
| | g.chr6:32610403C>A | 51348 | 14 | 0 | |
| | g.chr7:55238874T>A | 27957 | 6 | 0 | |
| | g.chr9:32459450T>G | 433 | 1 | 0 | |
| | 305 | | | | 926748 |
| | g.chr1:3428160T>G | 21762 | 27 | 0 | |
| | g.chr10:50944443T>A | 31031 | 21 | 0 | |
| | g.chr10:69959242C>A | 101233 | 41 | 0 | |
| | g.chr19:17330030C>A | 76859 | 54 | 0 | |
| | g.chr19:41383849C>G | 15509 | 18 | 0 | |
| | g.chr2:230861519G>T | 30579 | 17 | 0 | |
| | g.chr20:2290333C>A | 74840 | 45 | 0 | |
| | g.chr3:132166224C>A | 8383 | 12 | 0 | |
| | g.chr6:32610403C>A | 60275 | 36 | 0 | |
| | g.chr7:55238874T>A | 23397 | 11 | 0 | |
| | g.chr9:32459450T>G | 3462 | 6 | 0 | |
| | 396 | | | | 760326 |
| | g.chr1:3428160T>G | 13613 | 4 | 0 | |
| | g.chr10:50944443T>A | 27703 | 7 | 0 | |
| | g.chr10:69959242C>A | 65286 | 8 | 0 | |
| | g.chr19:17330030C>A | 171494 | 17 | 0 | |
| | g.chr19:41383849C>G | 0 | 0 | 0 | |
| | g.chr2:230861519G>T | 0 | 0 | 0 | |
| | g.chr20:2290333C>A | 42578 | 8 | 0 | |
| | g.chr3:132166224C>A | 0 | 0 | 0 | |
| | g.chr6:32610403C>A | 2599 | 2 | 0 | |
| | g.chr7:55238874T>A | 0 | 0 | 0 | |
| | g.chr9:32459450T>G | 10865 | 4 | 0 | |
| L107 | 128 | | | | 670825 |
| | g.chr1:152191717A>T | 84960 | 764 | 0 | |
| | g.chr10:69934258C>G | 20792 | 547 | 0 | |
| | g.chr11:128333503T>C | 55775 | 1131 | 0 | |

| | | | | | |
|---|---|---|---|---|---|
| | g.chr11:60102507G>T | 86798 | 862 | 0 | |
| | g.chr16:89212430C>T | 112304 | 618 | 0 | |
| | 213 | | | | 917898 |
| | g.chr1:152191717A>T | 132127 | 71 | 0 | |
| | g.chr10:69934258C>G | 44631 | 60 | 0 | |
| | g.chr11:128333503T>C | 49665 | 59 | 0 | |
| | g.chr11:60102507G>T | 192967 | 97 | 0 | |
| | g.chr16:89212430C>T | 203457 | 89 | 0 | |
| | 367 | | | | 743457 |
| | g.chr1:152191717A>T | 93722 | 68 | 0 | |
| | g.chr10:69934258C>G | 36753 | 62 | 0 | |
| | g.chr11:128333503T>C | 43708 | 66 | 0 | |
| | g.chr11:60102507G>T | 136302 | 89 | 0 | |
| | g.chr16:89212430C>T | 153579 | 85 | 0 | |
| L108 | 105 | | | | 1028891 |
| | g.chr1:11561278G>C | 75765 | 100 | 0 | |
| | g.chr1:57173271C>G | 58336 | 73 | 0 | |
| | g.chr11:36597104C>G | 34382 | 74 | 0 | |
| | g.chr13:58208018C>T | 75606 | 93 | 0 | |
| | g.chr16:30977694A>G | 99102 | 99 | 0 | |
| | g.chr17:17881032C>A | 19201 | 80 | 0 | |
| | g.chr17:6023871C>A | 1415 | 79 | 0 | |
| | g.chr2:33412052A>T | 29288 | 65 | 0 | |
| | g.chr20:19956215C>T | 67193 | 91 | 0 | |
| | g.chr6:31797631C>T | 56077 | 98 | 0 | |
| | g.chr7:117232188A>G | 13693 | 42 | 0 | |
| | g.chr8:22548872G>T | 76640 | 107 | 0 | |
| | g.chr9:135458585C>T | 269 | 46 | 0 | |
| | 208 | | | | 1024881 |
| | g.chr1:11561278G>C | 71282 | 135 | 0 | |
| | g.chr1:57173271C>G | 60311 | 102 | 0 | |
| | g.chr11:36597104C>G | 31496 | 94 | 0 | |
| | g.chr13:58208018C>T | 57732 | 108 | 0 | |
| | g.chr16:30977694A>G | 84436 | 127 | 0 | |
| | g.chr17:17881032C>A | 17957 | 111 | 0 | |
| | g.chr17:6023871C>A | 941 | 82 | 0 | |
| | g.chr2:33412052A>T | 27401 | 95 | 0 | |
| | g.chr20:19956215C>T | 65554 | 137 | 0 | |
| | g.chr6:31797631C>T | 59733 | 154 | 0 | |
| | g.chr7:117232188A>G | 7717 | 51 | 0 | |
| | g.chr8:22548872G>T | 50207 | 137 | 0 | |
| | g.chr9:135458585C>T | 191 | 33 | 0 | |

**Supplementary Table 10.** Read count data for bespoke cfDNA sequencing libraries from nine patients with localized prostate cancer, sampled from postoperative plasma. Reference and alternate reads count only unique molecules. Read depth: total number of mapped and unmapped reads.

**Supplementary Table 11**

| *Name* | *Sequence* |
|---|---|
| 502 | AATGATACGGCGACCACCGAGATCTACACCTCTCTATTCGTCGGCAGCGTCAGATGTGTATAAGAGACAG |
| 503 | AATGATACGGCGACCACCGAGATCTACACTATCCTCTTCGTCGGCAGCGTCAGATGTGTATAAGAGACAG |
| 504 | AATGATACGGCGACCACCGAGATCTACACAGAGTAGATCGTCGGCAGCGTCAGATGTGTATAAGAGACAG |
| 505 | AATGATACGGCGACCACCGAGATCTACACGTAAGGAGTCGTCGGCAGCGTCAGATGTGTATAAGAGACAG |
| 506 | AATGATACGGCGACCACCGAGATCTACACACTGCATATCGTCGGCAGCGTCAGATGTGTATAAGAGACAG |
| 507 | AATGATACGGCGACCACCGAGATCTACACAAGGAGTATCGTCGGCAGCGTCAGATGTGTATAAGAGACAG |
| 508 | AATGATACGGCGACCACCGAGATCTACACCTAAGCCTTCGTCGGCAGCGTCAGATGTGTATAAGAGACAG |
| 517 | AATGATACGGCGACCACCGAGATCTACACGCGTAAGATCGTCGGCAGCGTCAGATGTGTATAAGAGACAG |
| 701 | CAAGCAGAAGACGGCATACGAGATTCGCCTTAGTCTCGTGGGCTCGGAGATGTGTATAAGAGACAG |
| 702 | CAAGCAGAAGACGGCATACGAGATCTAGTACGGTCTCGTGGGCTCGGAGATGTGTATAAGAGACAG |
| 703 | CAAGCAGAAGACGGCATACGAGATTTCTGCCTGTCTCGTGGGCTCGGAGATGTGTATAAGAGACAG |
| 704 | CAAGCAGAAGACGGCATACGAGATGCTCAGGAGTCTCGTGGGCTCGGAGATGTGTATAAGAGACAG |
| 705 | CAAGCAGAAGACGGCATACGAGATAGGAGTCCGTCTCGTGGGCTCGGAGATGTGTATAAGAGACAG |
| 706 | CAAGCAGAAGACGGCATACGAGATCATGCCTAGTCTCGTGGGCTCGGAGATGTGTATAAGAGACAG |
| 707 | CAAGCAGAAGACGGCATACGAGATGTAGAGAGGTCTCGTGGGCTCGGAGATGTGTATAAGAGACAG |
| 708 | CAAGCAGAAGACGGCATACGAGATCCTCTCTGGTCTCGTGGGCTCGGAGATGTGTATAAGAGACAG |
| 709 | CAAGCAGAAGACGGCATACGAGATAGCGTAGCGTCTCGTGGGCTCGGAGATGTGTATAAGAGACAG |
| 710 | CAAGCAGAAGACGGCATACGAGATCAGCCTCGGTCTCGTGGGCTCGGAGATGTGTATAAGAGACAG |
| 711 | CAAGCAGAAGACGGCATACGAGATTGCCTCTTGTCTCGTGGGCTCGGAGATGTGTATAAGAGACAG |
| 712 | CAAGCAGAAGACGGCATACGAGATTCCTCTACGTCTCGTGGGCTCGGAGATGTGTATAAGAGACAG |

**Supplementary Table 11.** NGSO-4 oligonucleotide sequences ordered from Sigma in substitution of Illumina i5 and i7 adaptors. Oligonucleotides have longer regions of complementarity to primer sequences used for library preparation.

### SUPPLEMENTARY FIGURES

#### Supplementary Figure 1

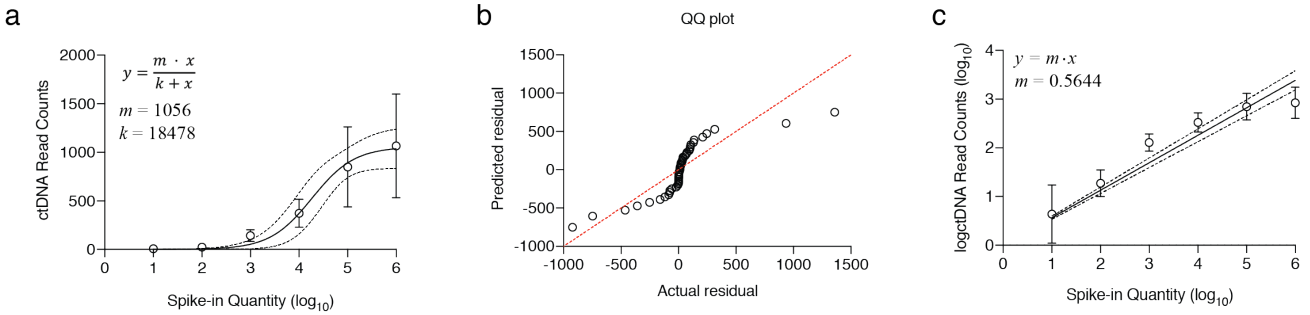


**Supplementary Figure 1. Recovery of mutant alleles from spike-in assays.** (a) Read counts of mutant alleles are plotted against the approximate spike in quantity and fitted to a nonlinear hyperbolic curve. Circles represent the average value across eight different targets, and error bars and dashed line depict the 95% confidence interval for the average and curve fit, respectively. (b) Quantile-quantile (QQ) plot of residuals from hyperbolic curve fit depicted in (a). Five outliers were detected but not removed from the plot. (c) Observed read counts were $\log_{10}$ transformed and plotted with linear regression.

**Supplementary Figure 2**

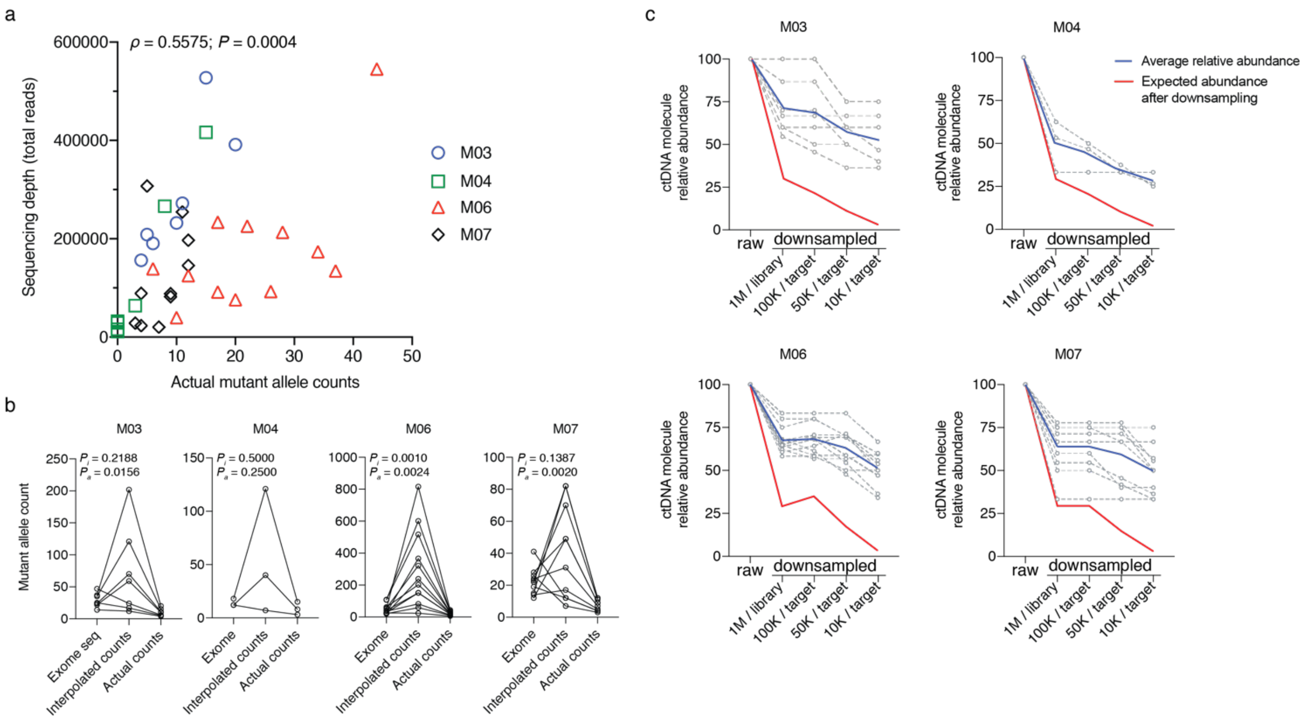


**Supplementary Figure 2. Performance of custom seq allele detection.** (a) Raw sequencing depth for each focus was plotted against actual mutant allele counts. Spearman's $\rho = 0.5575$; $P = 0.0004$. (b) Relationship between mutant allele counts from exome sequencing was compared against actual custom seq counts and adjusted (interpolated) counts. $P_i$: Wilcoxon matched-pairs signed rank test Exome seq vs. Interpolated counts. $P_a$: Wilcoxon matched-pairs signed rank test Exome seq vs. Actual counts. (c) Effect of downsampling on mutant allele detection by thresholds. Blue represents the observed average mutant allele count by performing downsampling prior to analysis. Red line represents the expected average mutant allele count performing downsampling after analysis.

#### Supplementary Figure 3

a

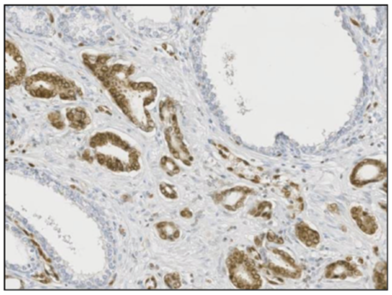

b

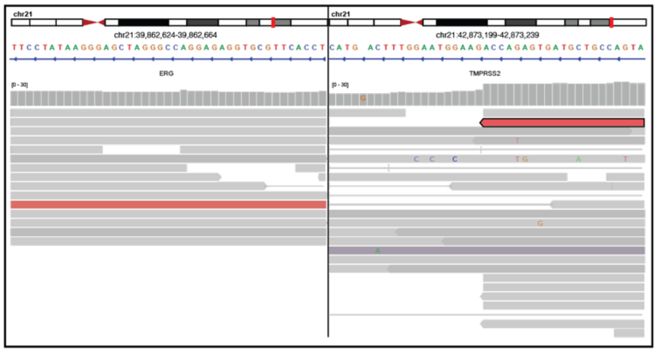

c

CTAAAGCTCCGGGCCACCCTCTCCCAGATGGGCCTCTTTCCACTTCCCCTTGCCAGCAGCTATA
TCTATGCCTCCCACCAGAGTGATGCTGCCAGTAACATTCTCCAGCTGGCCGGGCACGGCGGCT
CCTGCCTATAATTCCA

d

```
CTAAAGCTCCGGGCCACCCTCTCCCAGATGGGCCTCTTTCCACTTCCCCTTGCCAGCAGCTATA
                  -------------------->
                  forward outer primer   ------------------->
                                         forward nested primer

TCTATGCCTCCCACCAGAGTGATGCTGCCAGTAACATTCTCCAGCTGGCCGGGCACGGCGGCT
            ^                                                <------
       breakpoint                                            reverse
  <---- ERG | TMPRSS2 ---->    <--------------------
                               reverse nested primer

CCTGCCTATAATTCCA
---------------
outer primer
```

e

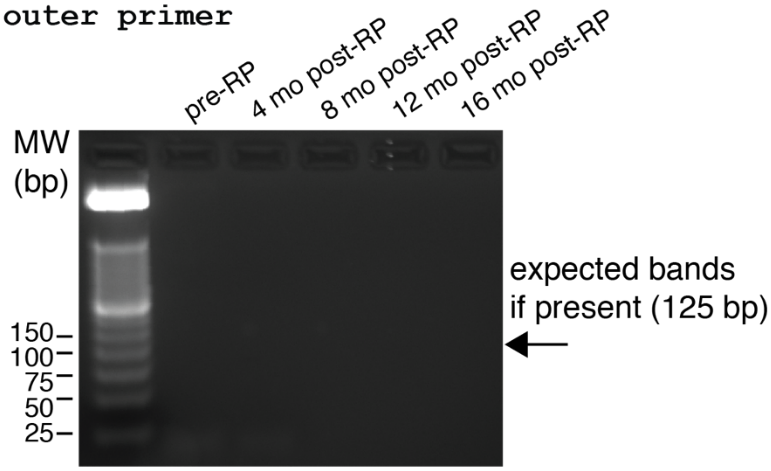


**Supplementary Figure 3. Detection of *TMPRSS2-ERG* breakpoint in ctDNA.** (a) anti-ERG immunohistochemistry of case L001 showing nuclear ERG overexpression in prostate cancer cells. (b) IGV screenshot showing WGS read (in red) mapping to chromosome 21 coordinates corresponding to *ERG* (left) and *TMPRSS2* (right). (c) BLAT results showing the breakpoint of the

WGS read. (d) Nested PCR strategy to amplify breakpoint from plasma cfDNA. (e) Photograph of agarose gel of PCR results.

#### Supplementary Figure 4

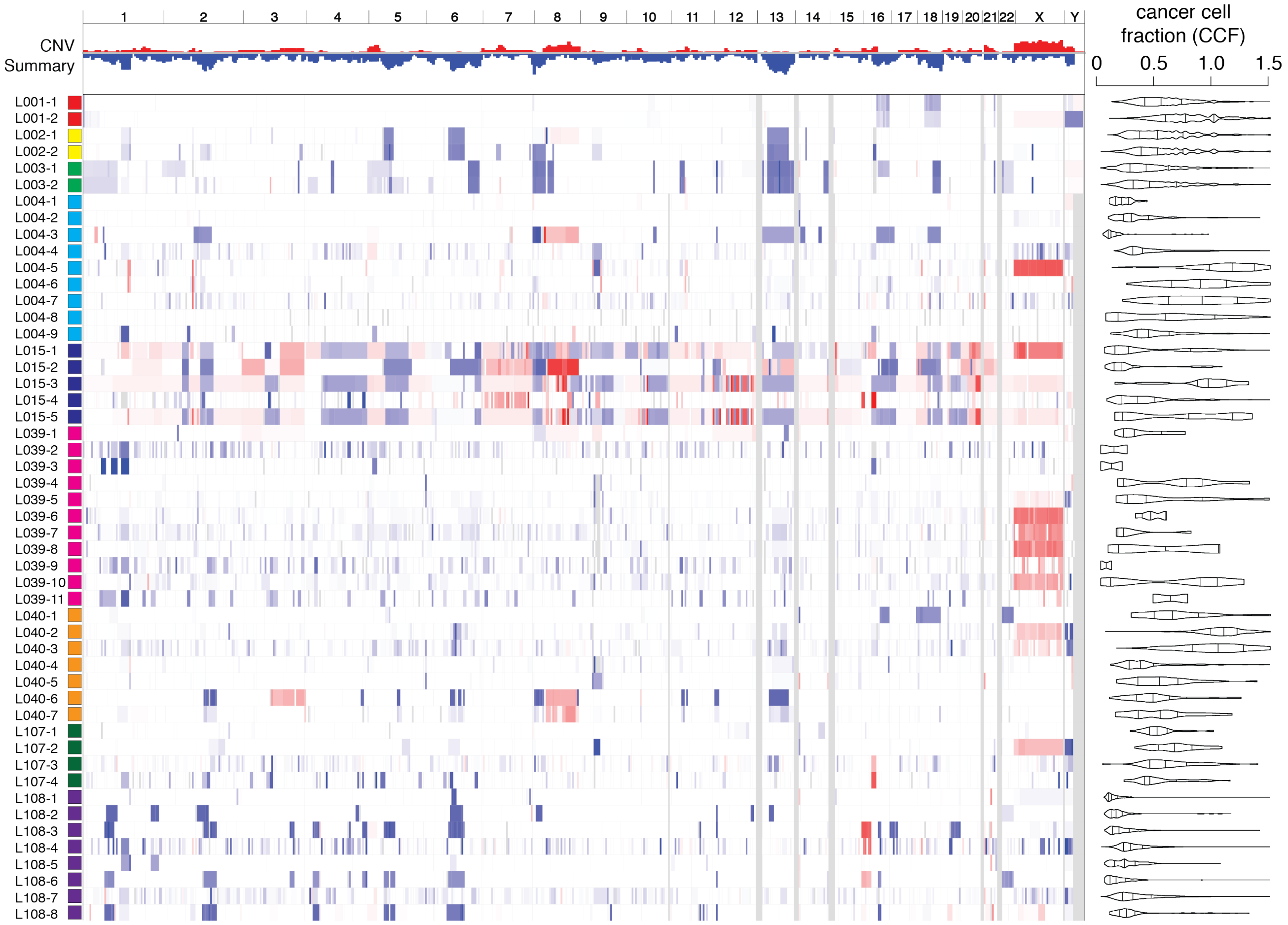


**Supplementary Figure 4. Genomic profile of localized prostate cancer.** For each laser capture microdissected focus (see Suppl. Table 5), somatic copy number alterations are shown on the left, integrated from read depth data from exome sequencing. The distribution of clonality measurements per point mutation, expressed as a cancer cell fraction (CCF), is shown on the right. Colored boxes on the left axis group each set of microdissected foci by patient.
